# Pateamine A mediates RNA sequence-selective translation repression by anchoring eIF4A and DDX3 to GNG motifs

**DOI:** 10.1101/2023.09.21.558742

**Authors:** Hironori Saito, Yuma Handa, Mingming Chen, Tilman Schneider-Poetsch, Yuichi Shichino, Mari Takahashi, Daniel Romo, Minoru Yoshida, Alois Fürstner, Takuhiro Ito, Kaori Fukuzawa, Shintaro Iwasaki

**Affiliations:** RNA Systems Biochemistry Laboratory, RIKEN Cluster for Pioneering Research, Wako, Saitama 351-0198, Japan; Department of Computational Biology and Medical Sciences, Graduate School of Frontier Sciences, The University of Tokyo, Kashiwa, Chiba 277-8561, Japan; School of Pharmacy and Pharmaceutical Sciences, Hoshi University, Shinagawa, Tokyo 142-8501, Japan; Chemical Genomics Research Group, RIKEN Center for Sustainable Resource Science, Wako, Saitama 351-0198, Japan; Laboratory for Translation Structural Biology, RIKEN Center for Biosystems Dynamics Research, Tsurumi-ku, Yokohama 230-0045, Japan; Department of Chemistry & Biochemistry, Baylor University, Waco, TX 76710, USA; Office of University Professor, The University of Tokyo, Bunkyo-ku, Tokyo 113-8657, Japan; Max-Planck-Institut für Kohlenforschung, D-45470 Mülheim/Ruhr, Germany; Graduate School of Pharmaceutical Sciences, Osaka University, Suita 565-0871 Japan

## Abstract

Small-molecule compounds that elicit mRNA-selective translation repression have attracted interest due to their potential for expansion of druggable space. However, only limited examples have been reported to date. Here, we show that pateamine A (PatA) represses translation in an mRNA-selective manner by clamping eIF4A, a DEAD-box RNA-binding protein, on GNG motifs. Through a systematic comparison of multiple eIF4A inhibitors by ribosome profiling, we found that PatA has unique mRNA selectivity in translation repression. Unbiased Bind-n-Seq revealed that PatA-targeted eIF4A exhibits a sequence preference for GNG motifs in an ATP-independent manner. This unusual RNA binding sterically hinders scanning by 40S ribosomes. *In silico* simulation, combination of classical molecular dynamics simulation and quantum chemical calculation, and the subsequent development of an inactive PatA derivative revealed that the positive charge of the tertiary amine on the trienyl arm induces G selectivity. Moreover, we identified DDX3, another DEAD-box protein, as an alternative target of PatA, showing the same effect as on eIF4A. Our results provide an example of the sequence-selective anchoring of RNA-binding proteins and mRNA-selective inhibition of protein synthesis by small-molecule compounds.

## Introduction

The production of harmful proteins often leads to deleterious outcomes in cells, causing a wide variety of diseases. Due to the limited druggable proteome ^1^, compounds that modulate the synthesis of unwelcome proteins at the translation level provide attractive therapeutic opportunities ^2^. Although several compounds that suppress translation in an mRNA-selective manner have been found ^2,3^, the number is still restricted, warranting the further identification of a new class with such activity.

Repurposing natural secondary metabolites for pharmacological use has been a common strategy in drug development ^4^. Indeed, translation inhibitors are not exceptions, as a variety of antibiotics targeting ribosomes have been exploited ^5^. In addition to ribosomes, eukaryotic translation initiation factor (eIF) 4A has been found to be a common target of a variety of natural products, presenting a vulnerability in cancer ^6^. These compounds include hippuristanol (Hipp) from a soft coral (*Isis hippuris*) ^7–11^, rocaglates from plants of the *Aglaia* genus ^12–18^, pateamine A (PatA) from a sponge (*Mycale* sp.) or its microbiome symbionts ^19–29^, and sanguinarine (San) from poppy plants (*Macleaya cordata* and *Argemone Mexicana*) ^30,31^.

eIF4A is an ATP-dependent DEAD-box type RNA-binding protein that forms a complex with cap-binding protein eIF4E and scaffold protein eIF4G and then facilitates the loading of the 43S preinitiation complex onto the 5′ ends of mRNA and subsequent scanning of the 5′ untranslated region (UTR) ^32,33^. In mammals, this protein is encoded in two genes, eIF4A1 and eIF4A2. Hipp and San have been shown to reduce the RNA-binding ability of eIF4A ^8–10,31^, simply inactivating the function of eIF4A in translation. In contrast, rocaglates have a unique mode of action, clamping eIF4A onto a polypurine (repeats of A and G nucleotides) RNA motif. This artificial clamping leads to steric hindrance to scanning ribosomes, blockage of the recruitement of 43S preinitiation complex at the 5′ ends of mRNAs, and ultimately the reudction of available eIF4A for translation initiation process ^15–18,34^. PatA also has been suggested not to phenocopy the loss of function of eIF4A, enhancing the interaction between eIF4A and RNA ^20–23,25,29^. However, the molecular mechanism of PatA-mediated protein synthesis blocking remained unclear.

Here, we systematically investigated the mode of action of PatA and found that this compound leads to RNA sequence-selective translation repression. Genome-wide ribosome profiling revealed that PatA induces distinct translational output compared to Hipp, San, and RocA—a potent rocaglate. RNA pulldown-Seq in cells and RNA Bind-n-Seq *in vitro* showed that PatA clamps on the GNG motif in RNA in an ATP-independent manner. PatA-mediated clamping causes mRNA-selective translation repression, most likely causing steric hindrance to scanning ribosomes. Our classical molecular dynamics simulation and subsequent quantum chemical calculation, revealed that the G nucleotide preference of PatA stemmed from the tertiary amine on the trienyl arm. The designed PatA derivative confirmed the importance of the amine in RNA selectivity, translation repression, and cytotoxicity. Our study provides an additional example of a sequence-selective translation inhibitor, expanding the space of exploitable proteomes for drug development.

## Results

### Differential impacts on cellular translation by multiple eIF4A inhibitors

To understand the variation in translation repression induced by eIF4A inhibitors and the mechanism of the effect evoked by PatA, we systematically compared the translatome alteration with ribosome profiling ^35,36^ (Figure 1A). Here, instead of the original PatA, we used the simplified derivative desmethyl desamino pateamine A (DMDA-PatA) (Figure S1A) due to their comparable activities ^37^. We conducted the experiments in HEK293 cell lines treated with Hipp, San, and DMDA-PatA at 2-3 different concentrations. To minimize the effect on the transcriptome, we treated cells with the compounds for 15-30 min, limiting the change in mRNA abundance ^15^. We also mined the published ribosome profiling data with RocA ^15^. Normalization by mitochondrial footprints as internal spike-ins ^17,34,38–40^ enabled us to monitor global translation changes. Indeed, the calculation of ribosome profiling data allowed us to monitor the dose-dependent translation alteration by the compounds (Figure S1B).

**Figure 1.**
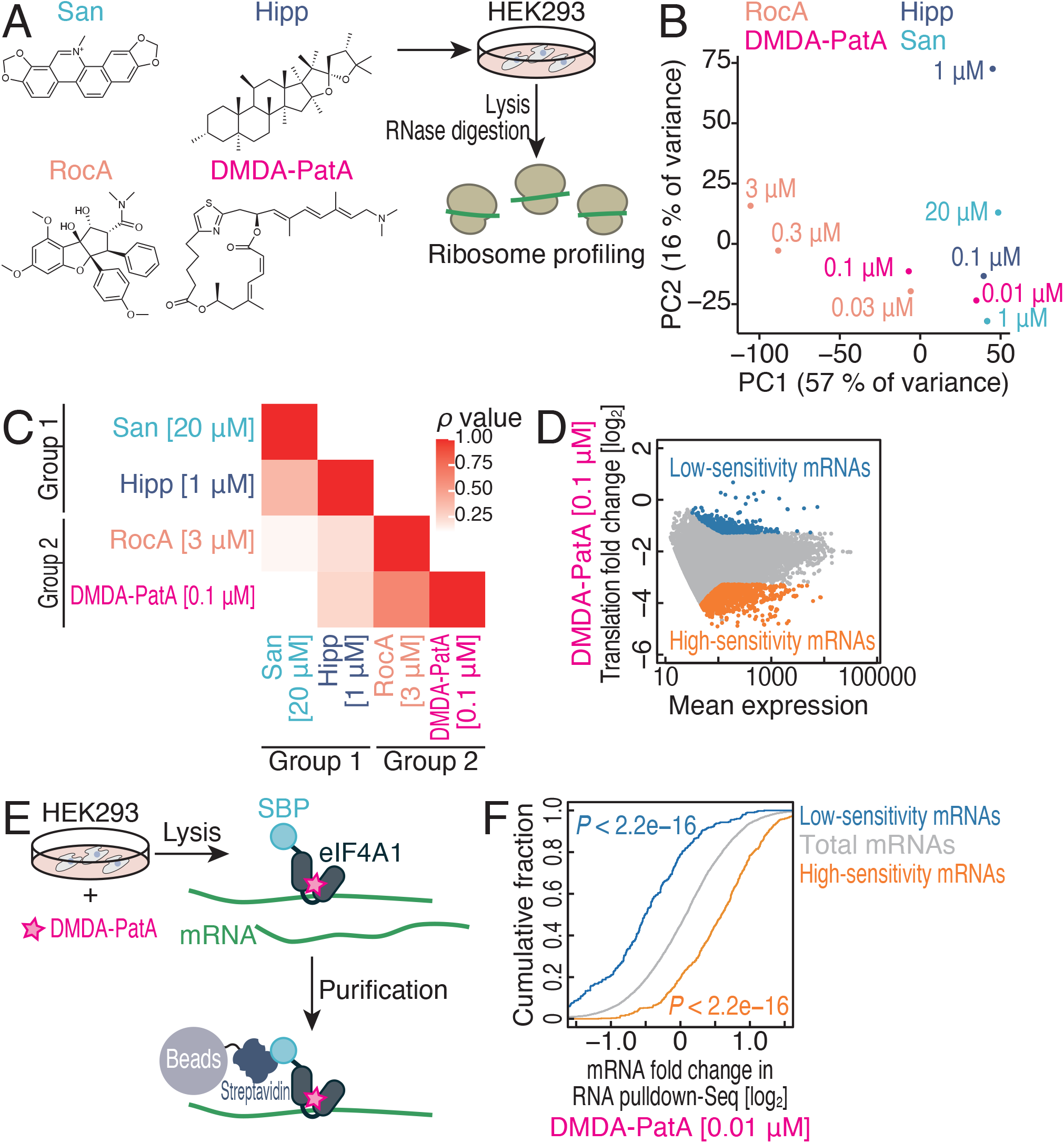
Comparative analysis of translation change by eIF4A targeting compounds in cells. (A) Schematic of ribosome profiling experiments. Chemical structures of eIF4A targeting compounds used in the experiments are shown. (B) Principal component analysis of the translation changes analyzed by ribosome profiling in the indicated conditions. (C) Spearman’s correlation coefficients (ρ) for translation changes induced by drug treatments. The color scales for ρ are shown. (D) MA (M, log ratio; A, mean average) plot of the translation fold change with 0.1 μM DMDA-PatA treatment. Low-sensitivity mRNAs (FDR ≤ 0.01 and log_2_-fold change ≥ 1 from the mean) and high-sensitivity mRNAs (FDR ≤ 0.01 and log_2_-fold change ≤ −1 from the mean) are highlighted. (E) Schematic of RNA pulldown-Seq experiments. mRNAs associated with SBP-tagged eIF4A1 in the cells were isolated and subjected to deep sequencing. (F) Cumulative distribution of the mRNA fold change in RNA pulldown-Seq for SBP-tagged eIF4A1 upon 0.01 μM DMDA-PatA treatment. DMDA-PatA low-sensitivity and high-sensitivity mRNAs (defined in D) were compared to total mRNAs. The significance was calculated by the Mann‒Whitney *U* test. See also Figure S1.

This comparative analysis revealed the similarities and differences in translation inhibition by eIF4A-targeting compounds. Principal component analysis (PCA) showed the dose dependent, directional effects of each drug (Figure 1B). Strikingly, eIF4A inhibitors could be categorized into two groups: group 1, San and Hipp; group 2, RocA and DMDA-PatA (Figures 1C and S1C-E).

Given that both San and Hipp reduce the affinity between eIF4A and RNA ^8,9,11,31,41^, their high correspondence in translation repression was a compelling result (Figures 1B-C and S1C-E). On the other hand, the polypurine-selective eIF4A clamping induced by RocA should be distinct from the effects of San and Hipp ^15–18,34^. Our analysis revealed that DMDA-PatA has a similar (but not identical) mode of translation repression to RocA, providing widespread sensitivity in translation repression across the transcriptome (Figure 1D).

### mRNA-selective clamping of eIF4A1 by DMDA-PatA leads to translation repression

Considering that PatA and its derivatives stabilize the interaction between RNA and eIF4A ^20–23,29^, we reasoned that the biased interaction of eIF4A with mRNAs may be associated with the mRNA selectivity of DMDA-PatA in translation repression. To monitor the mRNAs associated with eIF4A, we conducted RNA pulldown and subsequent RNA sequencing (RNA pulldown-Seq) with SBP-tagged eIF4A1, a major eIF4A paralog, from a HEK293 cell line (Figure 1E) ^15^. DMDA-PatA treatment evoked diverse alterations in mRNAs bound to eIF4A1 (Figure S1F).

Through comparison of the changes in mRNA association on eIF4A and translation repression, we found that the tight association between eIF4A and a subset of mRNAs upon drug treatment confers translation repression. mRNAs highly sensitive to DMDA-PatA in terms of translation repression were more stably associated with eIF4A upon drug treatment, whereas mRNAs with low sensitivity showed the opposite behavior (Figures 1F and S1G). These data suggested that the DMDA-PatA-mediated mRNA-selective eIF4A interaction determines the efficacy of translation repression.

### DMDA-PatA leads to ATP-independent GNG RNA clamping by eIF4A

These data led us to investigate whether DMDA-PatA clamps eIF4A on selective RNA motifs. To systematically survey the RNA motif selectivity provided by DMDA-PatA, we conducted RNA Bind-n-Seq ^42,43^ with random 30 nt RNA and recombinant human eIF4A1 in the presence and absence of DMDA-PatA (Figure 2A). eIF4A requires ATP to interact with RNA but dissociates from RNA upon ATP hydrolysis ^44,45^. To stabilize the ATP-bound ground state, we used a nonhydrolyzable ATP analog (5′-adenylyl-imidodiphosphate or AMP-PNP) for RNA Bind-n-Seq. Our analysis revealed the unexpected nucleotide specificity of DMDA-PatA toward a subset of motifs, which often included GNG sequences in 4-mer motif survey (Figure 2B). The same motif preference could be found when longer 5-mer or 6-mer motifs were considered (Figure S2A-B). Any nucleotide sandwiched by two Gs appeared be interchangeable in terms of selectivity (Figure 2C).

**Figure 2.**
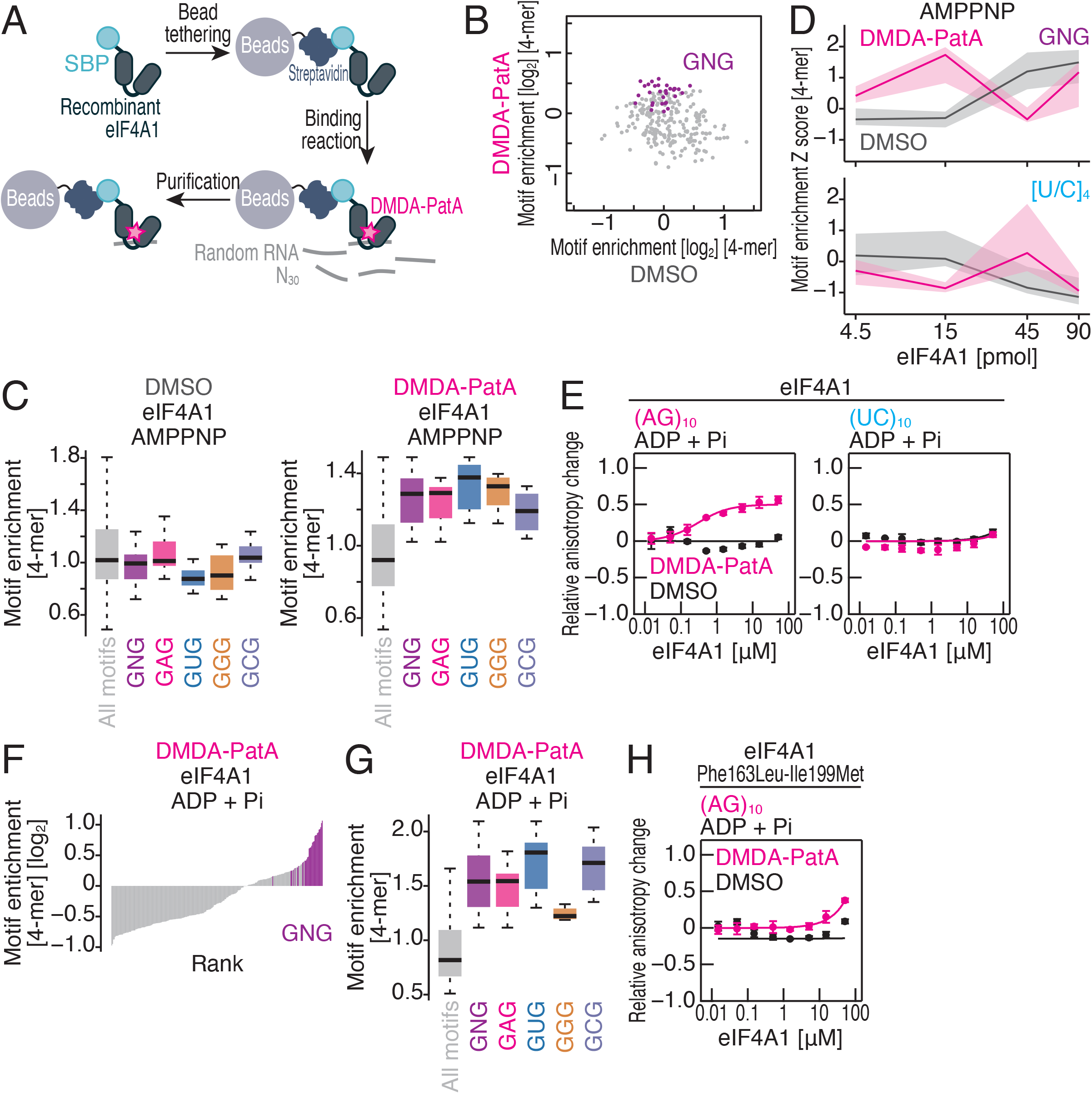
DMDA-PatA provides GNG motif preference on eIF4A1. (A) Schematic of RNA Bind-n-Seq experiments. Randomized RNAs associated with recombinant SBP-tagged eIF4A1 *in vitro* were isolated and subjected to deep sequencing. (B) Comparison of enrichment of 4-mer motifs with DMSO and those with DMDA-PatA. AMP-PNP and 15 pmol recombiannd eIF4A1 was included in the reaction. Motifs containing GNG are highlighted. (C) Box plots for motif enrichments in RNA Bind-n-Seq (with AMP-PNP) on eIF4A1 with DMSO or DMDA-PatA in the indicated 4-mer species. (D) Motif enrichment Z score along the titrated recombinant eIF4A1 in RNA Bind-n-Seq (with AMP-PNP) for the indicated 4-mer species with or without DMDA-PatA. (E and H) Fluorescence polarization assay for FAM-labeled RNAs along the titrated recombinant eIF4A1 (wild type or Phe163Leu-Ile199Met mutant) with ADP and Pi. The indicated RNA sequences at 10 nM were used with or without 50 μM DMDA-PatA. The data are presented as the mean (point) and s.d. (error) for replicates (n = 3). (F) Rank plot for 4-mer motifs enriched in RNA Bind-n-Seq (with ADP and Pi) on eIF4A1 in the presence of DMDA-PatA. Motifs containing GNG are highlighted. (G) Box plots for motif enrichments in RNA Bind-n-Seq (with ADP and Pi) on eIF4A1 with DMDA-PatA in the indicated 4-mer species. In box plots, the median (center line), upper/lower quartiles (box limits), and 1.5× interquartile range (whiskers) are shown. See also Figure S1.

To further monitor the affinity landscape of RNA-eIF4A1 binding, we titrated the amount of recombinant eIF4A1 for RNA Bind-n-Seq. With DMDA-PatA, the recovery of the GNG motif was increased at the middle amount (15 pmol) of eIF4A1 and then decreased at higher amounts of the protein (Figure 2D top). This peak in motif interaction could be expected since at high protein concentrations, the interaction with the strong-affinity motif became saturated, and competition was initiated with lower-affinity motifs^42^. In contrast, the interaction with irrelevant polypyrimidine sequences ([U/C]_4_) was enhanced at the higher concentration of eIF4A1 (at 45 pmol) by DMDA-PatA (Figure 2D bottom), indicating that this motif was a less preferred sequence for DMDA-PatA. A conventional fluorescence polarization assay with fluorescein (FAM)-conjugated RNAs confirmed these observations; GNG-possessing (AG)_10_ RNA associated more tightly with eIF4A1 upon DMDA-PatA treatment than the control (UC)_10_ RNA did. We note that DMDA-PatA enhanced the interaction of both RNAs with eIF4A1 (Figure S2C and Table 1), as reported in an earlier work ^29^.

**Table 1.** *K_d_* (μM) values between proteins and RNAs obtained in this study. The fluorescence polarization of FAM-labeled RNAs was determined and fitted to the Hill equation to determine *K_d_*. ND, not determined.

| eIF4A1 |  |  |  |  |  |
| --- | --- | --- | --- | --- | --- |
|  | ADP + Pi |  |  | AMP-PNP |  |
| RNA | DMSO | DMDA-PatA | iPr-DMDA-PatA | DMSO | DMDA-PatA |
| (AG)<sub>10</sub> | ND | $0.30 \pm 0.061$ | $26 \pm 17$ | $2.0 \pm 0.50$ | $0.41 \pm 0.053$ |
| (UC)<sub>10</sub> | ND | ND | | $28 \pm 5.1$ | $2.8 \pm 0.24$ |
| eIF4A1 Phe163Leu-Ile199Met |  |  |  |  |  |
|  | ADP + Pi |  |  | AMP-PNP |  |
| RNA | DMSO | DMDA-PatA | iPr-DMDA-PatA | DMSO | DMDA-PatA |
| (AG)<sub>10</sub> | ND | ND |  |  |  |
| eIF4A2 |  |  |  |  |  |

|  | ADP + Pi |  |  | AMP-PNP |  |
| --- | --- | --- | --- | --- | --- |
| RNA | DMSO | DMDA-PatA | iPr-DMDA-PatA | DMSO | DMDA-PatA |
| (AG)<sub>10</sub> | ND | 1.6 ± 0.17 |  |  |  |
| DDX3X (helicase core) |  |  |  |  |  |
|  | ADP + Pi |  |  | AMP-PNP |  |
| RNA | DMSO | DMDA-PatA | iPr-DMDA-PatA | DMSO | DMDA-PatA |
| (AG)<sub>10</sub> | ND | 0.80 ± 0.12 |  | 29 ± 61 | 0.71 ± 0.087 |
| (UC)<sub>10</sub> | ND | ND |  | ND | 1.3 ± 0.44 |
| DDX3X Gln360Leu (helicase core) |  |  |  |  |  |
|  | ADP + Pi |  |  | AMP-PNP |  |
| RNA | DMSO | DMDA-PatA | iPr-DMDA-PatA | DMSO | DMDA-PatA |
| (AG)<sub>10</sub> | ND | 5.1 ± 2.3 |  |  |  |
| DDX3X Gln360Pro (helicase core) |  |  |  |  |  |
|  | ADP + Pi |  |  | AMP-PNP |  |
| RNA | DMSO | DMDA-PatA | iPr-DMDA-PatA | DMSO | DMDA-PatA |
| (AG)<sub>10</sub> | ND | ND |  |  |  |

We found that the GNG-selective clamping of eIF4A1 could occur in an ATP-independent manner. In ADP and Pi, eIF4A1 *per se* could not bind to (AG)_10_ or (UC)_10_ RNA (Figure 2E and Table 1), as reported previously ^15^. However, DMDA-PatA enabled association with (AG)_10_ (Figure 2E and Table 1). This ATP-independent clamping did not occur on (UC)_10_ (Figure 2E and Table 1). To comprehensively survey the ATP-independent RNA selectivity, we again performed RNA Bind-n-Seq with ADP and Pi. As observed in the experiments with AMP-PNP, we detected a strong enrichment of GNG motifs (Figure 2F-G).

This ATP-independent sequence-selective clamping followed the reported biochemical modes of this compound. We observed that mutations in the binding interface of compound ^29^ (Phe163Leu-Ile199Met substitutions ^17^) attenuated the ATP-independent association of eIF4A1 with (AG)_10_ RNA in the presence of ADP and Pi (Figure 2H and Table 1). A PatA derivative was also known to target eIF4A2, a minor paralog of eIF4A ^29^. We found that ATP-independent GNG-selective clamping was also evoked on this paralog (Figure S2D and Table 1).

Thus, we conclude that DMDA-PatA provides GNG motif selectivity on eIF4A, evading the need for ATP.

### eIF4A clamping on the GNG motif sterically impedes scanning

We then investigated whether GNG motif-selective eIF4A clamping by DMDA-PatA could result in mRNA selectivity in translation repression in cells. Here, we calculated the correlation between motif numbers in the 5′ UTR and DMDA-PatA-mediated translational repression in ribosome profiling. Through a survey of all possible 4-mer motifs, we found more GNG motifs led to stronger repression by DMDA-PatA (Figure 3A). We note that in this analysis, all the motifs exhibited a negative correlation in general, probably due to the dependency of DMDA-PatA-mediated translation repression on 5′ UTR length (Figure S3A). Similarly, the GNG motif number was associated with tight mRNA interaction with eIF4A1 under DMDA-PatA treatment in RNA pulldown-Seq (Figure 3B). Thus, the presence of GNG motifs explained the selective translation repression by DMDA-PatA in cells.

**Figure 3.**
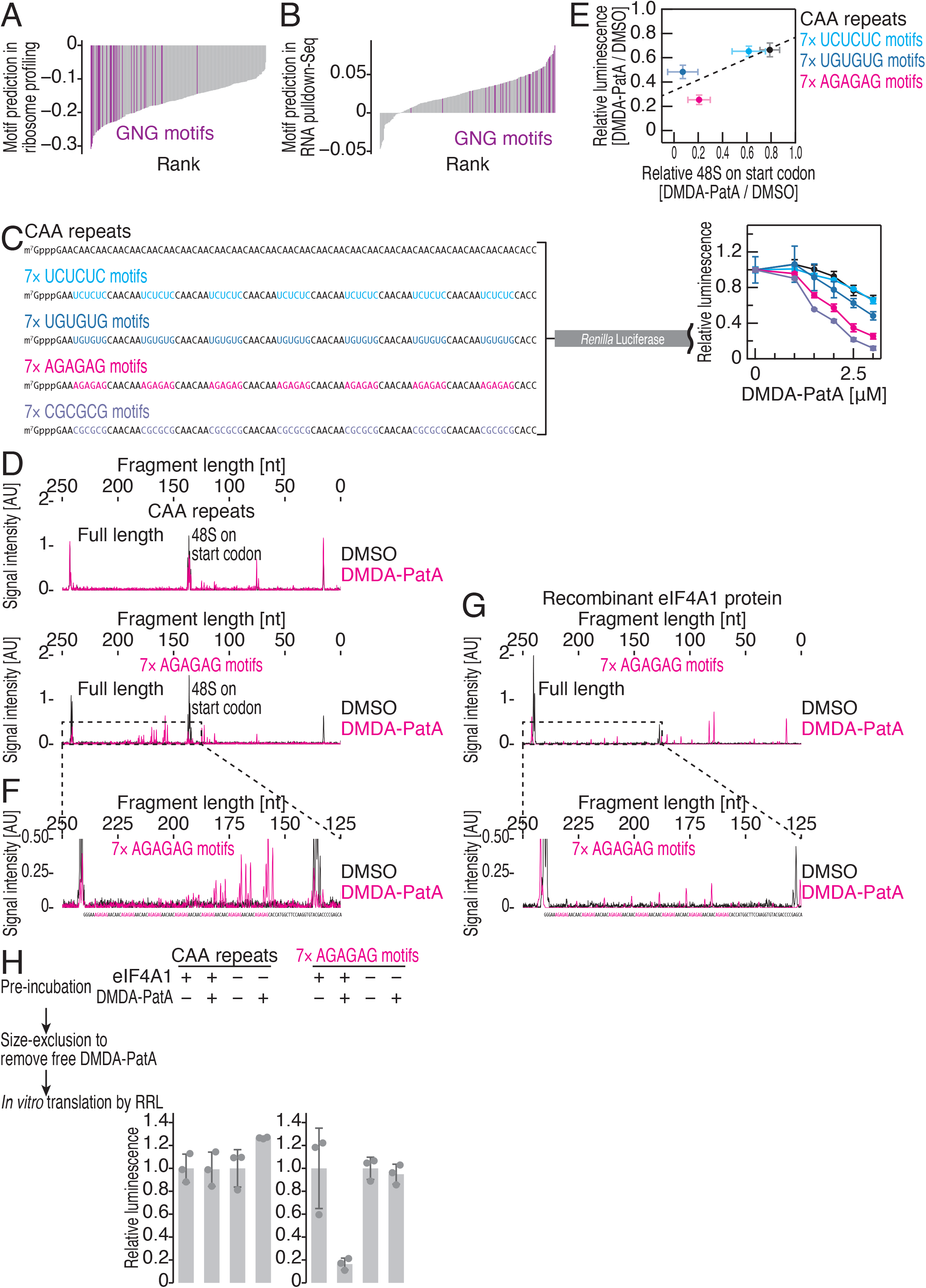
GNG motif-selective clamping of eIF4A causes the repression of translation initiation, related to Figure S3. (A) Rank plot for motif prediction by ribosome profiling in 0.1 μM DMDA-PatA treatment. Spearman’s correlation coefficients (ρ) between the number of 4-mer motifs found in 5′ UTRs and translation changes of the mRNAs were calculated. Motifs containing GNG are highlighted. (B) Rank plot for motif prediction by RNA pulldown-Seq in 0.01 μM DMDA-PatA treatment. Spearman’s correlation coefficients (ρ) between the number of 4-mer motifs found in 5′ UTRs and mRNA changes on SBP-tagged eIF4A1 were calculated. Motifs containing GNG are highlighted. (C) Schematic of reporter mRNAs with 7× NGNGNG motifs and the control CAA repeats (left). These mRNAs were subjected to *in vitro* translation with RRL and titration with DMDA-PatA. The data are presented as the mean (point) and s.d. (error) for replicates (n = 3). (D and F) Toeprinting assay to probe 48S ribosomes assembled on the start codons in the indicated reporter mRNAs with or without 10 μM DMDA-PatA. cDNA synthesized with FAM-labeled reverse transcription primers was analyzed by capillary electrophoresis. A magnified view of the results for reporter mRNA with 7× AGAGAG motifs (an area defined by dashed line in D) is shown in F. (E) The correspondence between translation repression observed in the *in vitro* translation (at 3 μM DMDA-PatA) (C) and the reduction of 48S formation (Figure S3E) for the indicated reporter mRNAs. The data are presented as the mean (point) and s.d. (error) for replicates (n = 3). The regression line (dashed line) is shown. (G) Toeprinting assay for recombinant eIF4A1 protein on the indicated reporter mRNAs with or without 10 μM DMDA-PatA. cDNA synthesized with FAM-labeled reverse transcription primers was analyzed by capillary electrophoresis. A magnified view of the results is shown at the bottom. (H) *In vitro* translation of reporter mRNA (with 7× AGAGAG motifs or CAA repeats, at 90.9 nM) preincubated with recombinant eIF4A1 and DMDA-PatA. The size exclusion column was used to eliminate free DMDA-PatA. The data are presented as the mean (bar) and s.d. (error) for replicates (point, n = 3). See also Figure S3.

We recapitulated the GNG motif-selective repression with an *in vitro* translation system using rabbit reticulocyte lysate (RRL). We prepared reporter mRNAs bearing unstructured CAA repeats ^46^ in the 5′ UTR as a control (Figure 3C). The substitution of a part of the CAA repeats with GNG motifs (7× UGUGUG, 7× AGAGAG, and 7× CGCGCG) (Figure 3C left) highly sensitized the reporter translation to DMDA-PatA (Figures 3C right and S3B). In contrast, the non-GNG motif 7× UCUCUC (Figure 3C left) did not affect DMDA-PatA sensitivity (Figures 3C and S3B). We observed motif number dependency of the translational repression (Figure S3C), as a single AGAGAG motif conferred weaker repression than 7 motifs (Figure 3C). We found limited positional effects on the single motif along the 5′ UTR (Figure S3C).

Using this setup, we further narrowed down the mechanism of DMDA-PatA-mediated translational repression. Here, we employed a toeprinting assay, which harnesses primer extension by reverse transcriptase and its blocking due to stable 48S formation on the start codon in the *in vitro* translation system ^15,46–48^. cDNAs extended with FAM-labeled primers were analyzed by capillary electrophoresis (Figures 3D, S3D, and S3E). This assay showed that DMDA-PatA suppresses 48S formation on AUG for GNG motif-containing reporters but not for the control CAA repeats (Figures 3D, S3D, and S3E). Overall, we found a correlation between translation repression efficiency and 48S formation blocking among the reporters (Figures 3E). These data indicated that DMDA-PatA inhibits translation initiation upstream of 48S formation on the start codon.

In our toeprinting assay on GNG motif-containing reporters, we detected additional cDNA fragments immediately downstream of GNG motifs upon DMDA-PatA treatment (Figure 3F), suggesting that the stable association of eIF4A in the lysate on these regions becomes a roadblock to reverse transcriptase. Indeed, recombinant eIF4A1 produced cDNA truncated downstream of GNG motifs, similar to that found in the lysate (Figure 3G).

This led us to test whether DMDA-PatA-mediated clamping on the 5′ UTR directly causes translational repression. For this purpose, the mRNA reporter preincubated with DMDA-PatA and recombinant eIF4A1 was subjected to *in vitro* translation with RRL ^15,17,49^. Purification through the gel-filtration column ensured the removal of free DMDA-PatA in the reaction. Nevertheless, protein synthesis from the reporter possessing GNG motifs was attenuated (Figures 3H and S3F). This experiment indicated that the clamped eIF4A by DMDA-PatA enabled the suppression of protein synthesis from the mRNA.

These results together revealed that RNA-selective eIF4A binding evoked by DMDA-PatA blocks translation, most likely by causing steric hindrance to scanning ribosomes.

### The tertiary amine on the trienyl arm confers GNG motif preference

The recent structure determination of the complex of desmethyl PatA (DM-PatA) (Figure S1A), eIF4A1, and polypurine RNA suggested that the compound does not have a clear interaction with RNA to discriminate bases ^29^. Thus, we wondered how PatA derivatives could exhibit sequence selectivity. To address this point, we performed classical molecular dynamics (MD) simulations and *ab initio* fragment molecular orbital (FMO) calculations ^50–53^ based on the reported structure of human eIF4A1•DM-PatA•polypurine RNA ^29^. For consistency with our experiments, we replaced DM-PatA with DMDA-PatA for the MD+FMO analysis ^54^.

To investigate the origin of the G preference, we replaced the RNA motif (_6_GAGA_9_) surrounding DMDA-PatA with _6_AGGA_9_, _6_AAAA_9_, or _6_GGGG_9_ (Figure 4A) and calculated inter-fragment interaction energy (IFIE) and its pair interaction energy decomposition analysis (PIEDA) ^55,56^ between those nucleotides and DMDA-PatA (Figure S4A). Among the 4 sequences analyzed, G nucleotides at positions 7 and 9 enabled more stable interactions with DMDA-PatA (Figure 4B-C). This stabilized interaction primarily stemmed from electrostatic energy terms (Figures 4D and S4A). These results provided an quantitative energetic explanation for the GNG motif preference conferred on eIF4A by DMDA-PatA.

**Figure 4.**
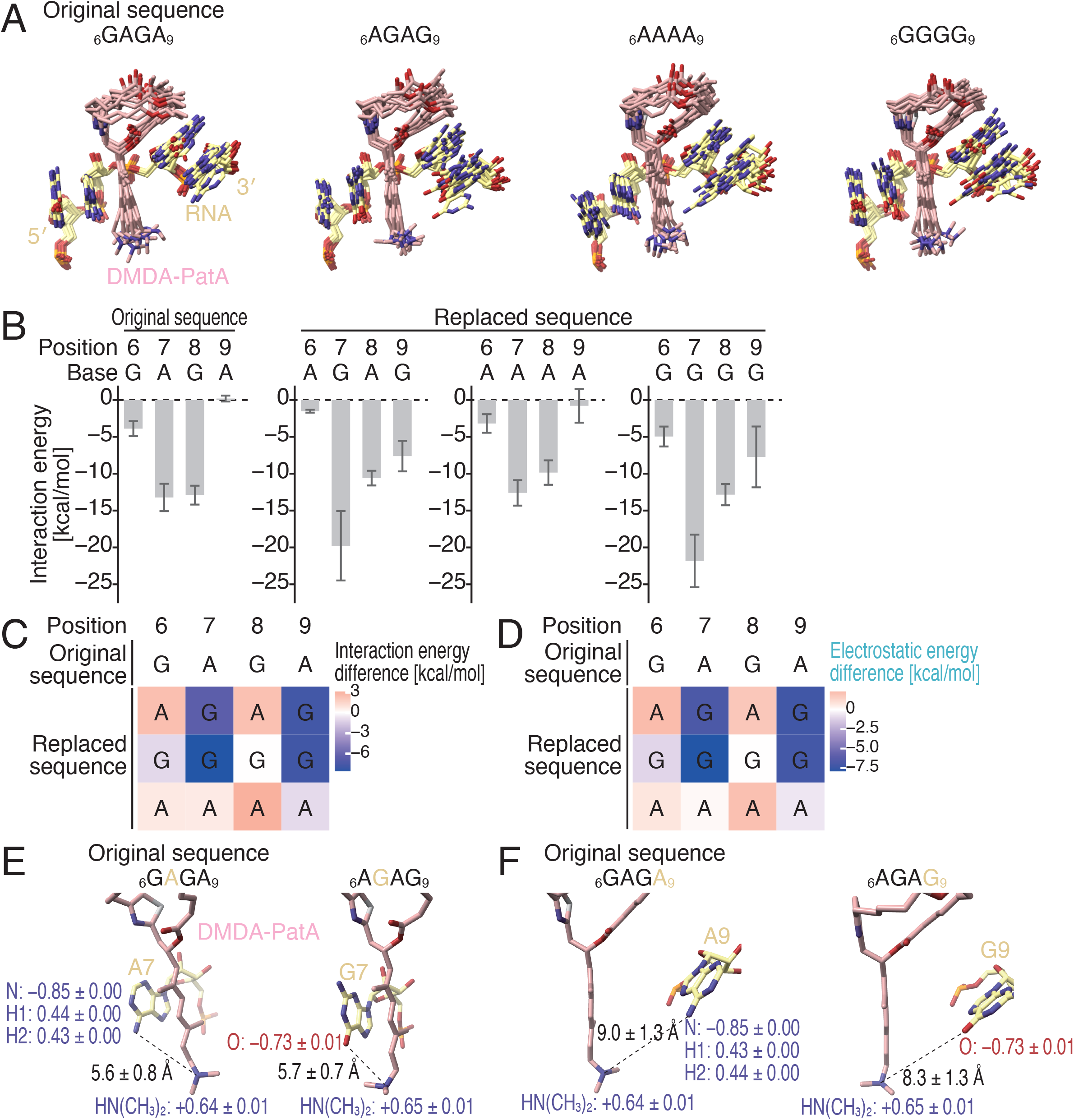
MD simulation and FMO calculation elucidated the energetic impact of RNA sequences on the association with DMDA-PatA. (A) Overlaid simulated structures (n = 10) of polypurine RNA•DMDA-PatA•eIF4A1 complexes analyzed by MD simulation. The complexes with RNAs possessing the indicated substitutions were investigated. (B) The interaction energy between DMDA-PatA and the indicated bases along the investigated complexes through FMO calculation. The data present the mean (bar) and s.d. (error) for 10 complexes. (C) The difference in the interaction energy between DMDA-PatA and the indicated bases from the original sequence (_6_GAGA_9_). See B for the source data. The color scale is shown. (D) The difference in the electrostatic energy between DMDA-PatA and the indicated bases from the original sequence (_6_GAGA_9_). See Figure S4A for the source data. The color scale is shown. (E and F) Representative structures obtained in MD simulation (at a time point of 100 ns) with _6_GAGA_9_ (original sequence) or _6_AGAG_9_. The net charge (*e*) on the indicated groups and the distances are shown (mean ± s.d. from 10 simulated structures). See also Figure S4.

We found that the tertiary amine on the trienyl arm of DMDA-PatA plays a key role in the electrostatic interaction with the G nucleotides at positions 7 and 9. MD+FMO analysis revealed that the tertiary amine was positively charged regardless of the RNA sequences (Figures 4E and S4B). This circumstance led to attractive interaction with the negatively charged O6 of the G nucleotide at position 7 (Figures 4E and S4B). On the other hand, the hydrogens on the amine at the same position in the A nucleotide did not result in charge attraction with the tertiary amine of DMDA-PatA (Figures 4E and S4B). We note that the similar distance from the tertiary amine of DMDA-PatA to the O6/N6 of the bases should not be the cause of base selectivity (Figures 4E and S4B). Essentially, similar theoretical explanation could be applied to the G and A nucleotides at position 9 (Figures 4F and S4C). These analyses provided an understanding for the G preference by DMDA-PatA on eIF4A1 at the atomic level.

### The tertiary amine of DMDA-PatA leads to sequence-selective translation repression

Given this observation, we directly tested whether a positively charged tertiary amine on the trienyl arm contributes to the G-rich motif preference. For this purpose, we prepared a PatA derivative in which the tertiary amine was replaced with an isopropyl group similar in size but nonbasic (Figures 5A and S5A, isopropyl-terminated DMDA-PatA or iPr-DMDA-PatA). Strikingly, RNA Bind-n-Seq (in the presence of ADP) with iPr-DMDA-PatA revealed that this compound no longer provided a GNG-motif preference (Figure 5B). The fluorescence polarization assay also supported this conclusion (Figure 5C, compared to Figure 2E, and Table 1). In contrast, other motifs were similarly recovered for both compounds, suggesting that iPr-DMDA-PatA only lost selectivity toward GNG motifs (Figure 5D). Due to the loss of sequence-selective eIF4A clamping, iPr-DMDA-PatA could not repress translation on reporter mRNAs with GNG motifs *in vitro* (Figure 5E, compared to Figure 3C). Moreover, global translation repression in cells and associated cell growth retardation were weakened in iPr-DMDA-PatA compared to DMDA-PatA (Figure 5F-G). We concluded that sequence-selective translation repression is caused by the tertiary amine on the trienyl arm of DMDA-PatA, directing cytotoxicity.

**Figure 5.**
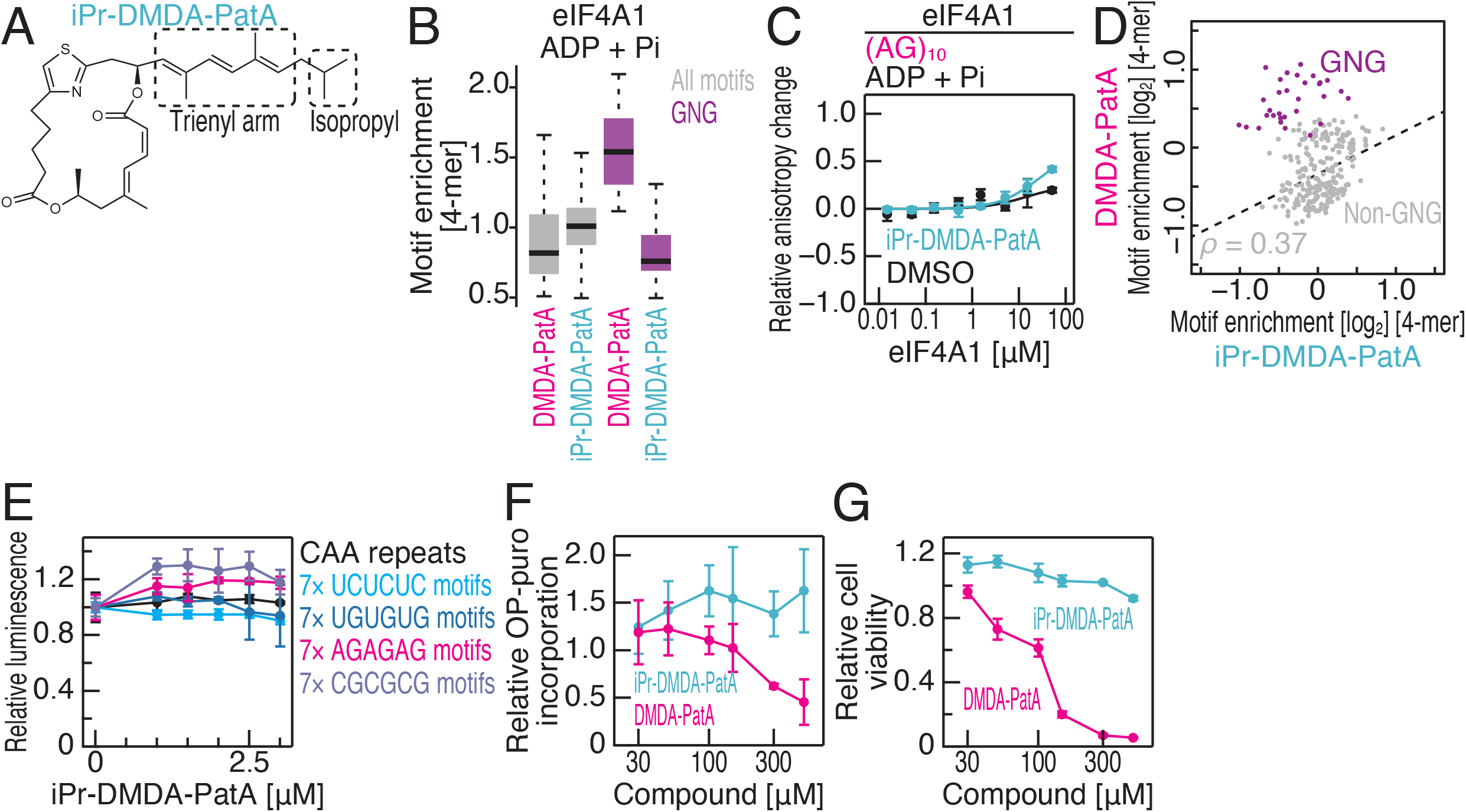
The tertiary amine on the trienyl arm of DMDA-PatA confers GNG selectivity on eIF4A1. (A) Chemical structures of iPr-DMDA-PatA. (B) Box plots for motif enrichments in RNA Bind-n-Seq (with ADP and Pi) on eIF4A1 with DMDA-PatA or iPr-DMDA-PatA in the indicated 4-mer species. (C) Fluorescence polarization assay for FAM-labeled RNAs along the titrated recombinant eIF4A1 with ADP and Pi. The indicated RNA sequences at 10 nM were used with or without 50 μM iPr-DMDA-PatA. The data are presented as the mean (point) and s.d. (error) for replicates (n = 3). (D) Comparison of enrichment of 4-mer motifs with DMDA-PatA and those with iPr-DMDA-PatA. ADP and Pi were included in the reaction. Motifs containing GNG are highlighted. The regression line (dashed line) for non-GNG motifs (gray points) is shown. ρ, Spearman’s correlation coefficients. (E) The mRNAs shown in Figure 3C left were subjected to *in vitro* translation with RRL with the titration of iPr-DMDA-PatA. The data are presented as the mean (point) and s.d. (error) for replicates (n = 3). (F) The relative OP-puro incorporation into nascent peptide in HEK293 cells was analyzed along the titrated DMDA-PatA or iPr-DMDA-PatA. The data are presented as the mean (point) and s.d. (error) for replicates (n = 3). (G) The relative cell viability was analyzed along the titrated DMDA-PatA and iPr-DMDA-PatA. The data are presented as the mean (point) and s.d. (error) for replicates (n = 3). See also Figure S5.

### Sequence selectivity differences between RocA and DMDA-PatA

Our results illuminated the similarity in translation repression mode between RocA and DMDA-PatA; both compounds clamp eIF4A on a subset of RNA motifs, presenting steric hindrance to scanning ribosomes ^15–18,34^. Although the translational impacts of these compounds were similar (Figure 1B-C), we noticed a substantial difference in the effects.

To profile the motif selectivity distinction of the compounds, we conducted RNA Bind-n-Seq experiments for RocA with titrated eIF4A1 recombinant protein in the presence of AMP-PNP. As reported previously ^15–18,34^, RocA enriched the polypurine ([A/G]_4_) sequences on eIF4A1 throughout the protein contents we tested (Figure S5B), rather than a peak in interaction at a specific protein amount. This suggested that the competition of less preferred motifs is limited in RocA. Considering the 3-mer motifs defined by polypurine and GNG motifs, we expected that GYG (where Y represents U or C) motifs, which were favorable for DMDA-PatA, would not be selected by RocA (Figure S5C). Consistent with this prediction, RNA Bind-n-Seq with RocA did not show enrichment in GYG motifs (Figure S5D). These data highlighted the similarities and differences in motif selection by the two compounds.

### DMDA-PatA also targets DDX3X for selective mRNA clamping and translation repression

Considering that RocA has been shown to target DDX3X ^34^, we were prompted to investigate the targeting potential of DMDA-PatA to DDX3X. We conducted a fluorescence polarization assay with recombinant DDX3X (helicase core) and observed its clamping on GNG motif-possessing (AG)_10_ RNA but not control (UC)_10_ RNA by DMDA-PatA, irrespective of the presence or absence of ATP (Figures 6A, S6A, and Table 1). More comprehensively, we conducted RNA Bind-n-Seq for recombinant DDX3X with ADP and found that DMDA-PatA allowed DDX3X to bind to GNG motifs (Figure 6B-C). Again, RocA and DMDA-PatA showed distinct sequence preferences on DDX3X as on eIF4A1 (Figure S6B).

**Figure 6.**
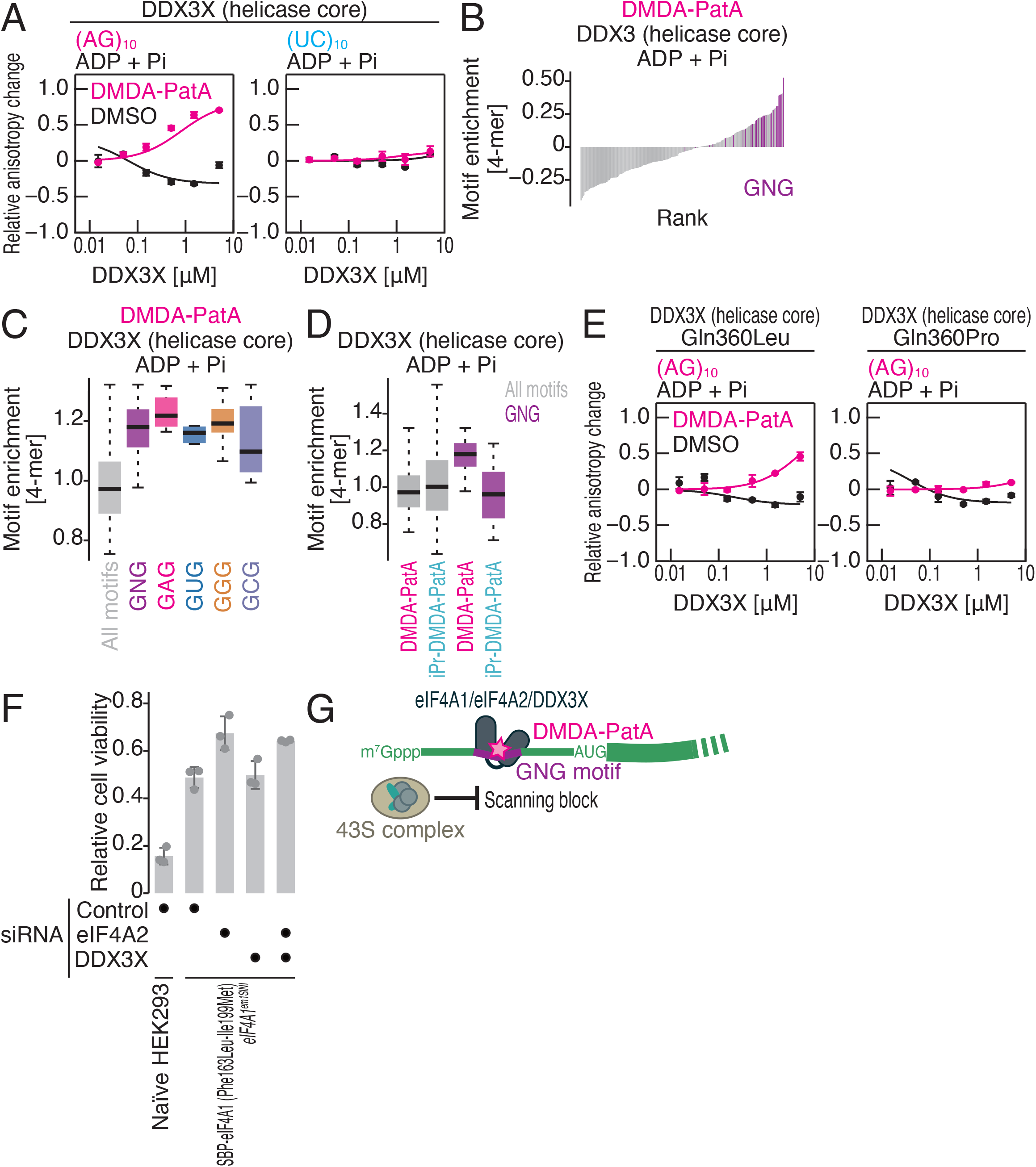
DMDA-PatA-mediated DDX3X clamping on the GNG motif in an ATP-independent manner, related to Figure S6. (A and E) Fluorescence polarization assay for FAM-labeled RNAs along the titrated recombinant DDX3X (helicase core) (wild type, Gln360Leu mutant, or Gln360Pro mutant) with ADP and Pi. The indicated RNA sequences at 10 nM were used with or without 50 μM DMDA-PatA. The data are presented as the mean (point) and s.d. (error) for replicates (n = 3). (B) Rank plot for 4-mer motifs enriched in RNA Bind-n-Seq (with ADP and Pi) on DDX3X (helicase core) in the presence of DMDA-PatA. Motifs containing GNG are highlighted. (B) Box plots for motif enrichments in RNA Bind-n-Seq (with ADP and Pi) on DDX3X (helicase core) with DMDA-PatA in the indicated 4-mer species. (C) Box plots for motif enrichments in RNA Bind-n-Seq (with ADP and Pi) on DDX3X (helicase core) with DMDA-PatA or iPr-DMDA-PatA in the indicated 4-mer species. (F) Relative cell viability for the indicated conditions after 0.1 μM DMDA-PatA treatment for 48 h. (G) Schematic of DMDA-PatA-mediated mRNA-selective translation repression. Clamping of eIF4A1/2 and/or DDX3X on the GNG motif in the 5′ UTR provides steric hindrance for scanning ribosomes.

This led us to test whether the RNA selectivity was also provided by the tertiary amine on the trienyl arm on DMDA-PatA. Indeed, RNA DDX3X Bind-n-Seq confirmed that iPr-DMDA-PatA lost the ability to confer GNG motif preference on the protein (Figure 6D), suggesting a base preference mechanism similar to that of eIF4A1-bound DMDA-PatA (Figure 4).

Considering that RocA depends on slightly different residues on DDX3X from eIF4A1 for binding ^34^, we investigated the role of those residues on DDX3X for DMDA-PatA targetability. As in the case of RocA ^34^, we confirmed the importance of Gln360 for DMDA-PatA-mediated ATP-independent clamping (Figure 6E and Table 1).

Finally, we tested the contributions of eIF4A1, eIF4A2, and DDX3X to DMDA-PatA-mediated cytotoxicity. Mutations in the RocA binding pocket (Phe163Leu-Ile199Met) ^17^ desensitized to DMDA-PatA in terms of cell viability (Figure 6F), consistent with the biochemical features (Figure 2H) ^29^. Additional knockdown of eIF4A2 further restored cell viability upon DMDA-PatA treatment (Figure 6F). In contrast, additional knockdown of DDX3X did not affect cytotoxicity, at least in the HEK293 cell lines that we used (Figure 6F). Given the DDX3X overexpression in a subset of cancer cells, the targeting of DMDA-PatA to this protein may be more significant in other cell types and should be considered for therapeutic purposes.

Taking these data together, we concluded that DMDA-PatA clamps eIF4A1, eIF4A2, and DDX3X on GNG RNA motifs on the 5′ UTR in an ATP-independent manner and presents steric hindrance to scanning ribosomes for mRNA-selective translation repression (Figure 6G).

## Discussion

Starting with the comparative study of eIF4A inhibitors, we found that PatA derivatives possessing the tertiary amine on the triene could elicit GNG motif preference by eIF4A1/2 and DDX3X DEAD-box RNA-binding proteins and inhibit protein synthesis from a subset of mRNAs. Our results provide an example of RNA-selective small molecules that have been hoped to unlock undruggable targets ^57,58^ but identified only in limited cases.

The importance of the tertiary amine on the trienyl arm has been revealed by the structure-activity relationship (SAR) analysis of PatA derivatives ^24^. However, the crystal structure of DM-PatA•human eIF4A1•polypurine RNA could not explain the significance of the terminal amine ^29^. Our work accounted for its role in RNA selectivity toward the GNG motif for eIF4A1/eIF4A2/DDX3X clamping, subsequently in mRNA-selective translation repression, and ultimately in cytotoxicity. Importantly, our work showed that PatA offers a unique framework of small molecules that can be used for RNA motif selection. This work will pave the way for the development of PatA derivatives for improved anticancer ^20,24,26^ and antiviral effects ^59–62^.

The recent structural analysis has showed that the binding pocket for PatA analog on eIF4A1 is largely overlapped with that for ocaglates ^29^. Although the modes to provide sequence selectivity are different, the sharply bent RNA, a characteristic conformation of eIF4A1-bound RNA, at the compound-binding interface ^17,29^ provides a unique context for sequence-selective clamping in both compounds. Our study examplified the convergence of two secondary metabolites distinct in their structures (Figure 1A) and their origins (marine sponge vs. plants) into a similar function in mRNA-selective translational repression.

## Acknowledgments

We thank all the members of the Iwasaki laboratory for constructive discussion and technical help. We are grateful to Dr. K. Dodo, Dr. K. Kato, Dr. C. Watanabe, and Dr. T. Honma for their helpful advice. Hipp was a kind gift from Dr. J. Tanaka. San was a kind gift from Dr. J. Liu. We thank Dr. C.-X. Zhuo and S. Schulthoff for the preparation of the pateamine derivatives and C. Wirtz for excellent NMR support (all at the MPI Mülheim). This study used the HOKUSAI SailingShip supercomputer facility at RIKEN; the Support Unit for Bio-Material Analysis, RIKEN CBS Research Resources Division; HiSeq 4000, supported by the National Institutes for Health (NIH) Instrumentation Grant (S10 OD018174), in QB3 Genomics, UC Berkeley, Berkeley, CA, (RRID:SCR_022170). MD simulations and FMO calculations were performed using the Fugaku supercomputer (project ID: hp220143) and the TSUBAME 3.0 supercomputer (Tokyo Institute of Technology, Japan). This work was supported by the Japan Society for the Promotion of Science (JSPS) [a Grant-in-Aid for Scientific Research (B), JP23H02415 (to S.I.); a Grant-in-Aid for Scientific Research (S), JP19H05640 (to M.Y.); a Grant-in-Aid for Scientific Research (C), JP23K05648 (to Y.S.)], the Ministry of Education, Culture, Sports, Science and Technology (MEXT) [a Grant-in-Aid for Transformative Research Areas (B), JP20H05784 (to S.I.); a Grant-in-Aid for Transformative Research Areas (A), JP21H05281 (to T.I.), JP21H05734, JP23H04268 (both to Y.S.); a Grant-in-Aid for Scientific Research on Innovative Areas, JP18H05503 (to M.Y.)], the Japan Agency for Medical Research and Development (AMED) [AMED-CREST, JP23gm1410001 (to S.I. and T.I.)], and RIKEN [Pioneering Projects “Biology of Intracellular Environments” (to S.I., T.I., and Y.S.); Incentive Research Projects (to T.S.-P.)]. This work was also supported by the Research Support Project for Life Science and Drug Discovery [Basis for Supporting Innovative Drug Discovery and Life Science Research (BINDS)] from AMED (JP23ama121030). H.S. was a RIKEN Junior Research Associate. A part of the *in sillico* study was conducted in an activity of FMO drug design consortium (FMODD) (to Y.H. and K.F.)

## Author contributions

Conceptualization, H.S. and S.I.;

Methodology, H.S., Y.H., M.C., T.S.-P., Y.S., M.T., and A.F.;

Formal analysis, H.S., Y.H., M.C., Y.S., M.T., and A.F.;

Investigation, H.S., H.S., Y.H., M.C., Y.S., M.T., and A.F.;

Resources, D.R. and A.F.;

Writing – Original Draft, S.I.;

Writing – Review & Editing, H.S., Y.H., M.C., T.S.-P., Y.S., M.T., D.R., M.Y., A.F., T.I., K.F., and S.I.;

Visualization, H.S. and S.I.;

Supervision, Y.S., M.Y., A.F., T.I., K.F., and S.I.;

Project administration, S.I.;

Funding Acquisition, T.S.-P., Y.S., M.Y., T.I., and S.I.

## Experimental procedures

### Compounds

RocA was purchased from Sigma‒Aldrich. Hipp and San were shared by Dr. Junichi Tanaka and Dr. Jun Liu, respectively. DMDA-PatA was synthesized in earlier studies ^37,63,64^. iPr-DMDA-PatA was synthesized as described below. These compounds were dissolved in dimethyl sulfoxide (DMSO).

### Chemical synthesis of iPr-DMDA-PatA

#### (Z)-4,4,5,5-Tetramethyl-2-(5-methylhex-2-en-2-yl)-1,3,2-dioxaborolane

The procedure was adapted from earlier work ^65^. [(ICy)CuCl] (ICy = N,N-dicyclohexylimidazolyl; 87.3 mg, 0.26 mmol) ^66^, NaO*t*-Bu (42.1 mg, 0.44 mmol) and B_2_pin_2_ (2.45 g, 9.63 mmol) were successively added to a solution of 5-methylhexan-2-one (1.00 g, 8.76 mmol), and the resulting mixture was stirred for 24 h at 70°C (bath temperature). *p*TsOH•H_2_O (3.33 g, 17.52 mmol) was then added, and stirring was continued for another 24 h at 65°C. After reaching ambient temperature, the suspension was filtered through a pad of Celite, which was carefully rinsed with CH_2_Cl_2_ in serval portions. The combined filtrates were concentrated under reduced pressure, and the crude material was purified by flash chromatography (hexane/*tert*-butyl methyl ether, 15:1) to give the title compound as a colorless oil (0.57 g, 29%). The analytical and spectroscopic data matched the literature ^67,68^.

#### iPr-DMDA-PatA

Pd(dppf)Cl_2_ (0.12 mg, 0.2 µmol) and Cs_2_CO_3_ (1.6 mg, 4.9 µmol) were successively added to a degassed solution of alkenyl iodide (1.8 mg, 3.2 µmol) ^64,69^ and alkenylpinacolboronate (0.73 mg, 3.2 µmol) in DMF (0.26 ml). The mixture was stirred overnight at ambient temperature and then diluted with *tert*-butyl methyl ether (0.5 ml) and washed with water (3 × 0.5 ml). The aqueous phase was extracted with CHCl_3_ (2 × 0.5 ml), and the combined organic layers were dried over Na_2_SO_4_. The solvent was evaporated under reduced pressure, and the residue was purified by flash chromatography (hexane/EtOAC, 10:1, complemented with 1% of Et_3_N) to give the title compound as a pale yellow solid (1.5 mg, 88%). [α]^20^_D_ = – 30.0 (*c* = 0.06, CHCl_3_). ^1^H NMR (C_6_D_6_, 600 MHz): δ 7.48 (dm, *J* = 11.6, 1.3 Hz, 1H), 6.73 (dddd, *J* = 9.6, 9.2, 4.5 Hz, 1H), 6.47 (t, *J* = 11.6 Hz, 1H), 6.36 (d, *J* = 15.9 Hz, 1H), 6.22 (d, *J* = 15.9 Hz, 1H), 6.19 (d, *J* = 1.0 Hz, 1H), 5.56 (d, *J* = 11.6 Hz, 1H), 5.51 (d, *J* = 9.6 Hz, 2H), 5.15 (dqd, *J* = 10.9, 6.4, 1.7 Hz, 1H), 3.08 (m, 2H), 2.79 (dtd, *J* = 14.5, 5.7, 4.3, 1.0 Hz, 1H), 2.46 (ddd, *J* = 16.10, 10.5, 6.4 Hz, 1H), 2.36 (ddd, *J* = 14.5, 10.5, 4.0, 1H), 2.14 (m, 1H), 2.09 (dd, *J* = 13.3, 10.9, 1H), 2.06 (m, 1H), 1.97 (d, *J* = 1.2 Hz, 2H), 1.97 (m, 3H), 1.72 (s, 3H), 1.64 (d, *J* = 13.3 Hz, 1H), 1.58 (m, 1H), 1.55 (s, 3H), 1.54–1.48 (m, 3H), 0.96 (d, *J* = 6.4 Hz, 3H), 0.88 (d, *J* = 6.7 Hz, 3H), 0.87 (d, *J* = 6.7 Hz, 3H). ^13^C NMR (C_6_D_6_, 151 MHz): δ 172.5, 165.4, 164.9, 157.3, 145.6, 141.2, 138.6, 134.9, 134.6, 133.1, 130.2, 128.7, 124.7, 115.5, 113.1, 69.9, 66.8, 48.4, 39.1, 37.9, 35.0, 31.2, 29.3, 28.5, 23.6, 22.6, 22.6, 21.1, 16.7, 13.4, 12.7. ^15^N NMR (C_6_D_6_, 61 MHz; via ^1^H-^15^N HMBC): δ -56.2. IR (film, cm^−1^): 2952, 2926, 2851, 1732, 1633, 1597, 1523, 1458, 1427, 1379, 1363, 1339, 1265, 1202, 1159, 1122, 1050, 1025, 986, 959, 814, 431. HRMS (ESI) *m/z* calcd. for C_31_H_43_NO_4_S+Na [*M*^+^ + Na]: 548.2807; found 548.2805.

### Ribosome profiling

The libraries used in this study are summarized in Table S1.

#### Library preparation

HEK293 cell lines were treated as follows: DMDA-PatA (0.01 μM or 0.1 μM) for 30 min, San (1 μM or 20 μM) for 30 min, and Hipp (0.1 μM or 1 μM) for 15 min. For the control, cells were incubated with 0.1% DMSO for the same durations as the drug treatments.

Library preparation was conducted following the protocol described previously ^70^. Cell lysates containing 10 μg of total RNA were subjected to RNase I (Lucigen) treatment for 45 min at 25°C. Ribosomes were collected using sucrose cushion ultracentrifugation. Subsequently, RNA fragments ranging from 26 to 34 nucleotides (nt) for Hipp treatment and from 17 to 34 nt for other samples were selected on a 15% denatured gel (FUJIFILM Wako Pure Chemical Corporation), followed by dephosphorylation and linker ligation. The removal of rRNA was performed utilizing the Ribo-Zero Gold rRNA Removal Kit (Illumina). The linker-conjugated RNAs were reverse-transcribed, circularized, and PCR-amplified. Single-end, 50-nt sequencing was performed utilizing HiSeq 4000 (Illumina).

#### Data analysis

Data were analyzed as previously reported ^71^. Briefly, the linker sequences were trimmed using fastx_clipper (http://hannonlab.cshl.edu/fastx_toolkit/index.html), followed by alignment of the reads to noncoding RNAs, including rRNAs, tRNAs, snoRNAs, snRNAs, and miRNAs, with Bowtie2 ^72^. Unaligned reads were mapped to the hg38 human genome reference and the custom mitochondrial transcript reference using Bowtie2. PCR duplicates were removed based on unique molecular identifiers (UMIs) on the linker sequences with a custom script (https://github.com/ingolia-lab/RiboSeq). The distance from the 5′ end to the ribosome A site on sequenced reads was empirically defined as follows: 15 for 26-30 nt, 16 for 31 nt, and 17 for 32 nt. The read count on each CDS was obtained with a custom script (https://github.com/ingolia-lab/RiboSeq), excluding the first and last 5 codons from the analysis. Regarding mitochondrial footprints, the A-site offset was set to 14 for 26-27 nt, 15 for 28-32 nt, 16 for 33-34 nt, and 17 for 35 nt. RocA-treated ribosome profiling data (0.03, 0.3, or 3 μM for 30 min) were published in an earlier work ^15^.

The change in ribosome footprint counts was calculated with DESeq ^73^. Subsequently, the data were renormalized to the average values of mitochondrial transcripts to calculate global translation change.

DMDA-PatA high-sensitivty mRNAs were defined as transcripts showing a log_2_-fold change of less than −1 from the mean with a false discovery rate (FDR) of less than 0.01 in 0.1 μM DMDA-PatA treatment. Conversely, low-sensitivity mRNAs were characterized as transcripts showing a log_2_ fold change of more than 1 from the mean with an FDR of less than 0.01.

Principal component analysis was conducted with a built-in function in R. Spearman correlations between the translation changes and the 4-mer numbers in the 5′ UTR were calculated to predict responsible motifs.

### RNA pulldown-Seq

The libraries used in this study are summarized in Table S1.

#### Library preparation

HEK293 Flp-In T-REx cells (Thermo Fisher Scientific, R78007) with SBP-tagged eIF4A1 integrant ^15^ were seeded in a 10-cm dish and cultured for 3 d in the presence of 1 μg/ml tetracycline. The cells were treated with 0.1% DMSO, 0.01 μM, or 0.1 μM DMDA-PatA for 30 min, washed with 5 ml of ice-cold PBS, and lysed with lysis buffer (20 mM Tris-HCl pH 7.5, 150 mM NaCl, 5 mM MgCl_2_, and 1 mM DTT) containing 1% Triton X-100 and 25 U/ml Turbo DNase (Thermo Fisher Scientific). The cell lysates were clarified by centrifugation at 20,000 × *g* for 10 min at 4°C. The supernatants were incubated with 30 μl of Dynabeads M-270 Streptavidin (Invitrogen), preequilibrated with lysis buffer containing 1% Triton X-100, for 30 min at 4°C. The beads were washed five times with lysis buffer containing 1% Triton X-100 and 1 M NaCl. The SBP-eIF4A1 and the bound RNAs were eluted from the beads by 40 μl of lysis buffer supplemented with 5 mM biotin for 30 min at 4°C. All buffers during the process above should contain 0.1% DMSO, 0.01 μM, or 0.1 μM DMDA-PatA. RNA was extracted with TRIzol LS reagent (Thermo Fisher Scientific) and the Direct-zol RNA MicroPrep Kit (Zymo Research). Sequencing libraries were generated utilizing a TruSeq Stranded Total RNA kit (Illumina) and sequenced on a HiSeq 4000 (Illumina) with 50-nt single-end reads.

#### Data analysis

Data were analyzed as previously reported ^15^. After the linker sequences were removed using fastx_clipper, the reads were aligned to ncRNAs such as rRNAs, tRNAs, snoRNAs, snRNAs, and miRNAs using STAR ^74^. Unaligned reads were mapped to the hg38 human genome reference using STAR.

The read counts for each transcript were obtained with the same custom script as described in the ribosome profiling section. Read fold change was calculated with DESeq. Spearman correlations between the mRNA changes and the 4-mer numbers in the 5′ UTR were calculated to predict responsible motifs.

### DNA construction

For His-tagged protein expression, pColdI-eIF4A1 WT ^17^, pColdI-eIF4A1 Phe163Leu-Ile199Met ^17^, pColdI-DDX3X helicase core WT ^34^, pColdI-DDX3X helicase core Gln360Leu ^34^, and pColdI-DDX3X helicase core Gln360Pro ^34^ were used. For His- and SBP-tagged protein expression, pColdI-SBP-eIF4A1 ^34^ and pColdI-SBP-DDX3X helicase core ^34^ were used.

### Recombinant protein purification

*E. coli* BL21 Star (DE3) cells (Thermo Fisher Scientific) transformed with the pColdI plasmids were cultured in 1 l of LB medium supplemented with ampicillin at 37°C to OD_600_ 0.6. Then, the cells were chilled for 30 min at 4°C, followed by overnight cultivation at 15°C in the presence of 1 mM IPTG. The cells were harvested by centrifugation at 8,000 × *g* for 2 min, flash-frozen in liquid nitrogen, and stored at −80°C.

The pellet was suspended in bacterial lysis buffer (20 mM HEPES, 500 mM NaCl, 10 mM imidazole, 0.5% NP-40, and 10 mM β-mercaptoethanol, adjusted to pH 7.5 by NaOH) and subsequently sonicated on ice. The lysate was clarified by centrifugation at 10,000 × *g* for 20 min at 4°C. The supernatant was then incubated with 3 ml of Ni-NTA Superflow agarose beads (QIAGEN), which were preequilibrated with bacterial lysis buffer, for 1 h at 4°C in a sealed gravity column (Bio-Rad). The beads on the gravity column were washed with 50 ml of high-salt wash buffer (20 mM HEPES, 1 M NaCl, 20 mM imidazole, and 10 mM β-mercaptoethanol, adjusted to pH 7.5 by NaOH) and then with 50 ml of low-salt wash buffer (20 mM HEPES-NaOH pH 7.5, 10 mM NaCl, 20 mM imidazole, and 10 mM β-mercaptoethanol, adjusted to pH 7.5 by NaOH). The His-tagged protein was eluted with 8 ml of elution buffer (20 mM HEPES, 10 mM NaCl, 250 mM imidazole, 10% glycerol, and 10 mM β-mercaptoethanol, adjusted to pH 7.5 by NaOH).

The eluted protein was further purified using an NGC chromatography system (Bio-Rad). Specifically, the protein was loaded on a HiTrap 1 ml Heparin HP column (Cytiva) and fractionated through a gradient of increasing salt concentration using a mixture of buffer A (20 mM HEPES-NaOH pH 7.5, 10 mM NaCl, 10% glycerol, and 1 mM DTT) and B (20 mM HEPES-NaOH pH 7.5, 1 M NaCl, 10% glycerol, and 1 mM DTT). The fractions containing the target protein were collected and buffer-exchanged to storage buffer (20 mM HEPES-NaOH pH 7.5, 150 mM NaCl, 10% glycerol, and 1 mM DTT) with a PD-10 column (Cytiva). The protein was concentrated using Amicon Ultra-4 10 kDa MWCO (Millipore) according to the manufacturer’s instructions. The recombinant protein was flash-frozen with liquid nitrogen and stored at −80°C.

### Fluorescence polarization assay

The reaction mixtures (10 μl each) were prepared as follows: 0-50 μM recombinant eIF4A1 or 0-5 μM recombinant DDX3X, 10 nM FAM-labeled RNA (Hokkaido System Science), 1% DMSO, 50 μM DMDA-PatA, or 50 μM iPr-DMDA-PatA, 1 mM AMP-PNP, 1mM MgCl_2_, 20 mM HEPES-NaOH pH 7.5, 150 mM NaCl, 1 mM DTT, and 5% glycerol. The mixtures were incubated at room temperature for 30 min and transferred to black 384-well microplates (Corning). Then, the anisotropy change was measured by an Infinite F-200 PRO (Tecan).

To test the ATP requirement, AMP-PNP was replaced with 1 mM ADP and 1 mM Na_2_HPO_4_.

Data were fitted to Hill equations with Igor Pro 8 (Wavemetrix) to estimate *K_d_*.

### RNA Bind-n-Seq

The libraries prepared in this study are summarized in Table S1.

#### Library preparation

For the experiment with AMP-PNP, 4.5, 15, 45, or 90 pmol of SBP-tagged eIF4A1 protein was incubated with 30 μl of Dynabeads M-270 Streptavidin (Thermo Fisher Scientific), which had been preequilibrated with equilibration buffer (20 mM Tris-HCl pH 7.5, 150 mM NaCl, 5 mM MgCl_2_, and 1 mM DTT) containing 1% Triton X-100, for 30 min at 4°C. The beads were washed 3 times with 60 μl of equilibration buffer containing 1% Triton X-100 and 1 M NaCl and then twice with 60 μl of equilibration buffer containing 0.1% Triton X-100. Subsequently, the beads were incubated with a 1 μM N_30_ oligonucleotide [5′-ctctttccctacacgacgctcttccgatct-N_30_-atcgtagatcggaagagcacacgtctgaa-3′ (Gene Design), where the lower cases represent the DNA sequence and N represents a random RNA sequence], for 30 min at 37°C in 30 μl of equilibration buffer containing 0.1% Triton X-100, 0.33 U/μl SUPERase•In RNase Inhibitor (Thermo Fisher Scientific), 2 mM AMP-PNP, and 3 μM DMDA-PatA. Then, the beads were washed 3 times with equilibration buffer containing 0.1% Triton X-100, 2 mM AMP-PNP, and 3 μM DMDA-PatA. The protein-oligonucleotide complexes bound to the beads were eluted using 30 μl of equilibration buffer containing 0.1% Triton X-100, 5 mM biotin, 2 mM AMP-PNP, and 3 μM DMDA-PatA for 30 min at 4°C. The oligonucleotides were purified with an Oligo Clean & Concentrator Kit (Zymo Research) and converted into a DNA library as described in the ribosome profiling section ^70^. For the control experiment, DMDA-PatA was substituted with the same volume of DMSO (0.1% in the reaction). For the input experiment, the random oligonucleotides were directly converted to a DNA library. Experiments with RocA were also conducted as described above but substituting 3 μM DMDA-PatA with 3 μM RocA.

For the experiments with ADP and Pi, experiments were performed with 2 mM ADP and 2 mM Na_2_HPO_4_ instead of 2 mM AMP-PNP. For complex assembly, 50 μM N_30_ oligonucleotide and 90 pmol of SBP-tagged recombinant proteins (eIF4A1 or DDX3X) were used. iPr-DMDA-PatA was also used at 3 μM throughout the experiments.

DNA libraries were sequenced on HiSeq 4000 (Illumina) with 50-nt single-end read mode or HiSeqX Ten (Illumina) with 150-nt paired-end mode.

#### Data analysis

In the case of paired-end reads, fastp ^75^ was employed to correct read errors, and read 1 was used for the downstream analysis. fastx_clipper was utilized to eliminate the linker sequence, followed by fastx_collapser, which aggregated identical sequences into single sequences.

The frequency of all possible 4-mers was calculated, and the motif enrichment was expressed as the ratio to that in the input library.

RNA Bind-n-Seq with RocA in the presence of ADP and Pi was obtained from an earlier work ^34^.

### *In vitro* translation

The reaction mixture (10 μl) was prepared with 5 μl of rabbit reticulocyte lysate nuclease-treated (Promega), 2 μl of H_2_O, 1 μl of DMDA-PatA or iPr-DMDA-PatA dissolved in 1% DMSO, 1 μl of 500 nM mRNA reporter, and 1 μl of premix [100 μM amino acid mixture minus methionine (Promega), 100 μM amino acid mixture minus leucine (Promega), and 0.5 U/μl SUPERase•In RNase Inhibitor (Thermo Fisher Scientific)] and incubated for 1 h at 30°C. After the translation reaction was quenched by adding 30 μl of 1× Passive Lysis Buffer (Promega), 10 μl of the mixture was placed on a 96-well white assay plate (Coster), and the fluorescence signal was measured by the *Renilla*-Glo Luciferase Assay System (Promega) and GloMax Navigator System (Promega).

To perform *in vitro* translation for preformed mRNA reporter complexes, a reaction (27.5 μl) containing 9.1 μM recombinant eIF4A1, 9.1 μM DMDA-PatA (dissolved in 2% DMSO), 90.9 nM reporter mRNA with 7× AGAGAG motifs or CAA repeats, 16.6 mM HEPES-NaOH at pH 7.5, 55.3 mM potassium acetate, 2.8 mM magnesium acetate, 1.8 mM ATP, and 552.7 μM DTT was incubated for 5 min at 30°C. Subsequently, 2.5 μl of 285 nM magnesium acetate was added to the mixture. The reaction mixture was loaded into a MicroSpin G-25 column (Cytiva) that had been equilibrated with buffer containing 30 mM HEPES-NaOH at pH 7.5, 100 mM potassium acetate, 1 mM magnesium acetate, and 1 mM DTT and centrifuged at 700 × *g* for 1 min at 4°C to eliminate free DMDA-PatA. The eluted fraction was mixed with 2.5 μl of storage buffer (20 mM HEPES-NaOH at pH 7.5, 150 mM NaCl, 10% glycerol, and 1 mM DTT). Then, 4 μl of the eluted solution was combined with 5 μl of RRL and 1 μl of premix and incubated for 1 h at 30°C. In the case of experiments using mRNA reporters with 7× CGCGCG motifs, the concentrations of mRNA reporters were adjusted to 181.8 nM.

In the control experiments, the recombinant eIF4A1 protein was replaced with the storage buffer in the preformation reaction, and the recombinant eIF4A1 protein was added to the G-25 column flowthrough instead of the storage buffer. DMDA-PatA was substituted with 2% DMSO.

### Toeprinting assay

A reaction consisting of 0.5× RRL, 2 mM GMPPNP, 2.5 mM magnesium acetate, and 10 μM DMDA-PatA (with 0.2% DMSO) in 10 μl was incubated for 5 min at 30°C. Then, after the addition of 1 μl of 500 nM mRNA reporter, the mixture was further incubated for 5 min at 30°C. Subsequently, the resulting mixture was combined with 9 μl of RT mix [22.2 mM Tris-HCl pH 7.5, 111.1 mM KCl, 3.2 mM magnesium acetate, 1.1 mM DTT, 1.1 mM each dNTP (New England Biolabs), 27.8 nM 5′ fluorescein-labeled primer (5′-/56FAM/ATGCAGAAAAATCACGGC-3′, eurofins), and 22.2 U/μl ProtoScript II Reverse Transcriptase (New England Biolabs)] and incubated for 15 min at 30°C. cDNAs were purified using a Direct-zol RNA MicroPrep kit with TRIzol LS reagent (Thermo Fisher Scientific). Subsequently, the cDNAs were subjected to a second purification with AMPure XP beads (Beckman Coulter). In the control experiments, the reaction contained 0.2% DMSO instead of DMDA-PatA. Instead of 0.5× RRL and 2 mM GMPPNP, 2.5 μM recombinant eIF4A1 protein and 2 mM ATP were also used.

The purified cDNAs were analyzed with GeneScan 400HD ROX dye Size Standard (Thermo Fisher Scientific) on an Applied Biosystems 3130*xl* Genetic Analyzer (Thermo Fisher Scientific). The data were analyzed using Peak Scanner 2.

Dideoxy-terminator sequencing was employed to calibrate the GeneScan 400HD ROX dye Size Standard for the cDNA length synthesized in the current experimental setups. The reaction containing 25 nM mRNA reporter, 12.5 nM RT primer, 0.5 mM (each) dNTPs, 0.5 mM ddNTP (ddATP, ddTTP, ddGTP, or ddCTP), ProtoScrip II Reverse Transcriptase, and 1× ProtoScript II RT Reaction Buffer (New England Biolabs) was incubated for 1 h at 30°C. cDNAs were purified with an Oligo Clean & Concentrator Kit and analyzed as described above.

### MD and FMO calculations

The complex structure of the DM-PatA•eIF4A1•polypurine RNA complex was obtained from the Protein Data Bank (PDB) (6XKI) ^29^. DMDA-PatA was created by removing amines from DM-PatA using the Molecular Operating Environment (MOE, https://www.chemcomp.com/Products.htm). Subsequently, hydrogen atoms not determined by X-ray crystallography were added using the “Protonate 3D” function in MOE, considering a protonation state at pH 7.0. Afterward, the atomic coordinates were optimized. Moreover, the DNA/RNA builder of MOE was utilized to generate complexes in which the _6_GAGA_9_ sequences in the RNA surrounding DMDA-PatA were substituted with _6_AGGA_9_, _6_AAAA_9_, and _6_GGGG_9_. All modeling in MOE was performed using the AMBER10:EHT force field.

MD simulations were performed for the four complexes created for 50 ns. A heat process from 0 K to 310 K was performed for 50 ps using the NVT ensemble. Next, an equalization process was performed at 310 K for 50 ps (NPT ensemble). Furthermore, density relaxation was performed for 1 ns (NPT ensemble), and a production run was performed for 100 ns at 310 K (NPT ensemble). Note that the pressure at NPT was 1013 hPa. The force fields used in this MD simulation were Amberff14SB for the protein, OL3 for the RNA, and Gaff2 for the DMDA-PatA. TIP3P water was utilized as the solvent, and Na^+^ ions were used as the counterions. The bond distances involving hydrogen were not constrained. The time step was 1 fs. This MD simulation was conducted under periodic boundary conditions. Furthermore, the MD simulations in this study were executed using the AMBER16 program (https://ambermd.org/doc12/Amber16.pdf).

From the 50-ns trajectories obtained in the MD simulations, we extracted 10 structures at 3 ns intervals starting from 73 ns, resulting in a total of 40 structures. The geometry of each sampled structure was optimized by applying constraints on the heavy atoms. Then, FMO calculations were performed. The ABINIT-MP program (<u>Tanaka et al. 2014</u>; <u>Mochizuki et al. 2021</u>) was used for the FMO calculations; electron correlation effects were incorporated by second-order Møller–Plesset perturbation (MP2) theory, which was efficiently implemented in ABINIT-MP ^76–78^. For the basis functions ^52,53^, we used 6-31G*, a standard of FMO calculations. Subsequently, the average value and standard deviation of 40 structures of total IFIE with eIF4A and RNA for DMDA-PatA obtained using these FMO calculations were calculated for each pose. In addition, PIEDA divides the IFIE into four energy components, *i.e.*, electrostatic (ES), exchange repulsion (EX), charge transfer (CT), and dispersion (DI), allowing the physicochemical properties of molecular interactions to be evaluated.

### Cell viability assay

On 24-well plates, 500 μl of 4 × 10^4^ cells/ml HEK293 Flp-In T-REx cells (Thermo Fisher Scientific, R78007) or HEK293 Flp-In T-REx SBP-eIF4A1 (Phe163Leu-Ile199Met) *eIF4A1^em1SINI^* cells ^17^ was seeded and incubated overnight. Transfection was performed with 55 nM DDX3X-specific siRNA (Dharmacon, L-006874-02-0005) and eIF4A2-specific siRNA (Dharmacon, L-013758-01-0005) using the TransIT-X2 Transfection Reagent System (Mirus). Following 2 d of incubation, 200 μl of 2 × 10^4^ cells/ml cells were seeded into individual wells of a 96-well plate and incubated for 6 h. The siRNA knockdown was repeated once more, with the same protocol above. After 24 h of additional incubation, the cells were subjected to treatment with 0.1 μM DMDA-PatA (with 0.1% DMSO) or just 0.1% DMSO for 48 h. The cell viability was assessed utilizing the RealTime-Glo MT Cell Viability Assay System (Promega). The luminescence was measured using a GloMax Navigator System (Promega).

For the experiments without transfection, 4,000 HEK293 Flp-In T-REx cells were seeded on a 96-well plate and treated with different concentrations of DMDA-PatA or iPr-DMDA PatA for 24 h. Cell viability was assessed as described above.

### Nascent peptide labeling with OP-puro

On 24-well plates, 500 μl of 5 x 10^5^ cells/ml HEK293 Flp-In T-REx cells were seeded and then incubated for 24 h. Subsequently, 50 μl of 0.22 mM *O*-propargyl-puromycin (OP-puro, Jena Bioscience) dissolved in Opti-MEM (Thermo Fisher Scientific) was added to the medium with varying concentrations of DMDA-PatA or iPr-DMDA-PatA. The cells were incubated for 30 min, washed with PBS, and then lysed with an OP-puro lysis buffer (20 mM Tris-HCl pH 7.5, 150 mM NaCl, 5 mM MgCl_2_, and 1% Triton X-100). The lysate was centrifuged at 20,000 × *g* for 10 min at 4°C. The supernatant was subjected to a click reaction with IRdye800CW azide (LI-COR) following the protocol provided in the Click-iT Cell Reaction Buffer Kit (Thermo Fisher Scientific). The reaction mixture was loaded onto a MicroSpin G-25 Column (Cytiva), which was equilibrated with an OP-puro lysis buffer containing 1 mM DTT and centrifuged at 700 × *g* for 2 min. After the proteins were separated with SDS‒PAGE, the infrared 800 nm (IR800) signal on the gel was detected using Odyssey CLx (LI-COR). Subsequently, the SDS‒PAGE gel was stained with Coomassie brilliant blue (CBB) (EzStainAQua, ATTO), and the total protein abundance was monitored by Odyssey CLx with the IR700 channel. OP-puro incorporation (IR800 signal) was normalized to the total protein abundance (IR700 signal).

## Data and code availability

The results of ribosome profiling, RNA pulldown-Seq, and RNA Bind-n-Seq (GEO: GSE243312) obtained in this study have been deposited in the National Center for Biotechnology Information (NCBI) database. All input and result files for FMO calculations are available at the FMODB (https://drugdesign.riken.jp/FMODB/) (see Table S2 for the ID list).

---

**Figure S1.**
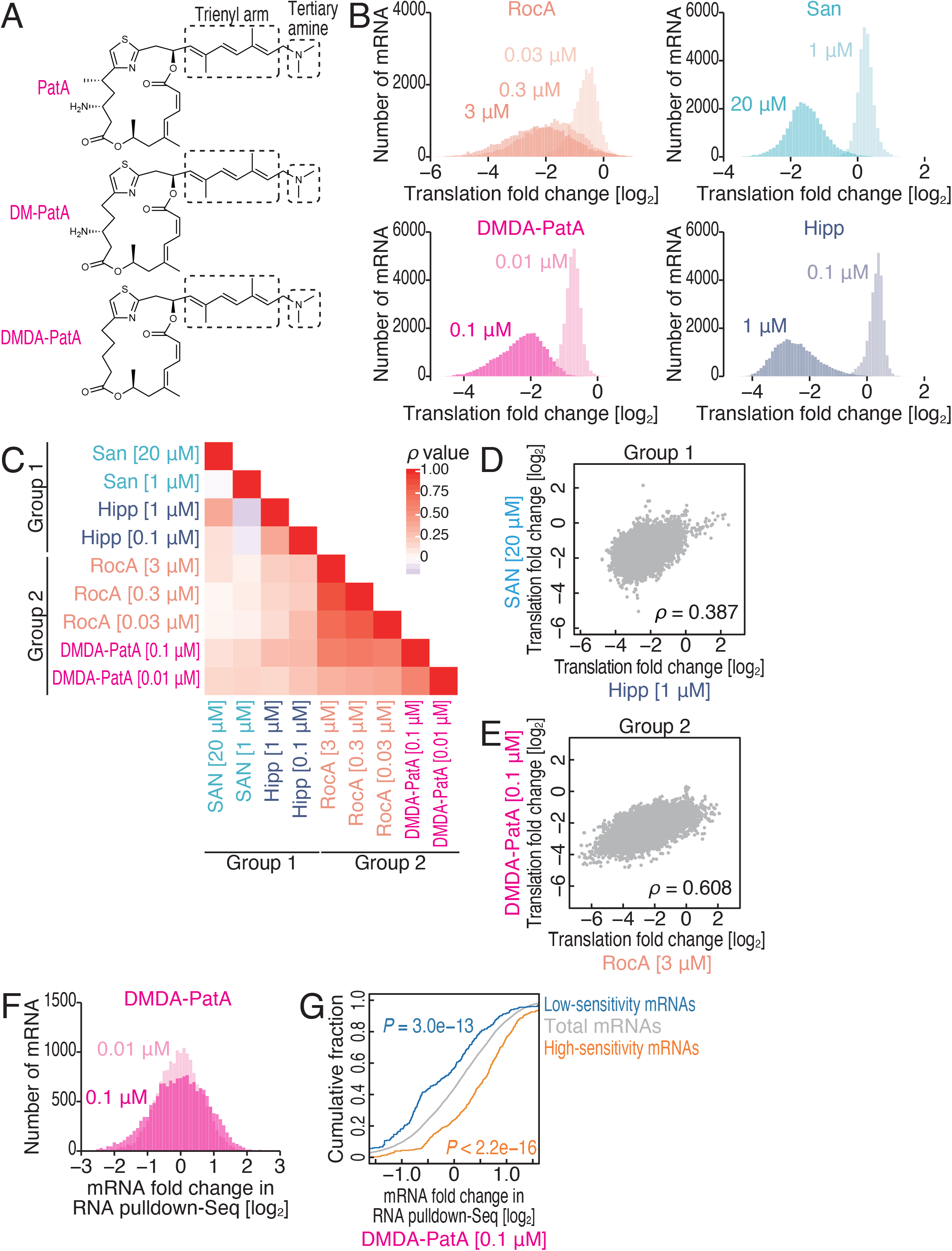
Characterization of ribosome profiling and RNA pulldown-Seq with eIF4A inhibitors, related to Figure 1. (A) Chemical structures of PatA derivatives. (B) Histograms of translation changes measured by ribosome profiling under the indicated conditions. (C) Spearman’s correlation coefficients (ρ) of translation changes induced by drug treatments. The color scales for ρ are shown. (D and E) Comparison of ribosome profiling data in group 1 compounds (D) and in group 2 compounds (E). ρ, Spearman’s correlation coefficient. (F) Histograms of mRNA changes recovered in SBP-eIF4A1 measured by RNA pulldown-Seq under the indicated conditions. (G) Cumulative distribution of the mRNA fold change in RNA pulldown-Seq for SBP-tagged eIF4A1 upon 0.1 μM. DMDA-PatA low-sensitivity and high-sensitivity mRNAs (defined in Figure 1D) were compared to total mRNAs. The significance was calculated by the Mann‒Whitney *U* test.

**Figure S2.**
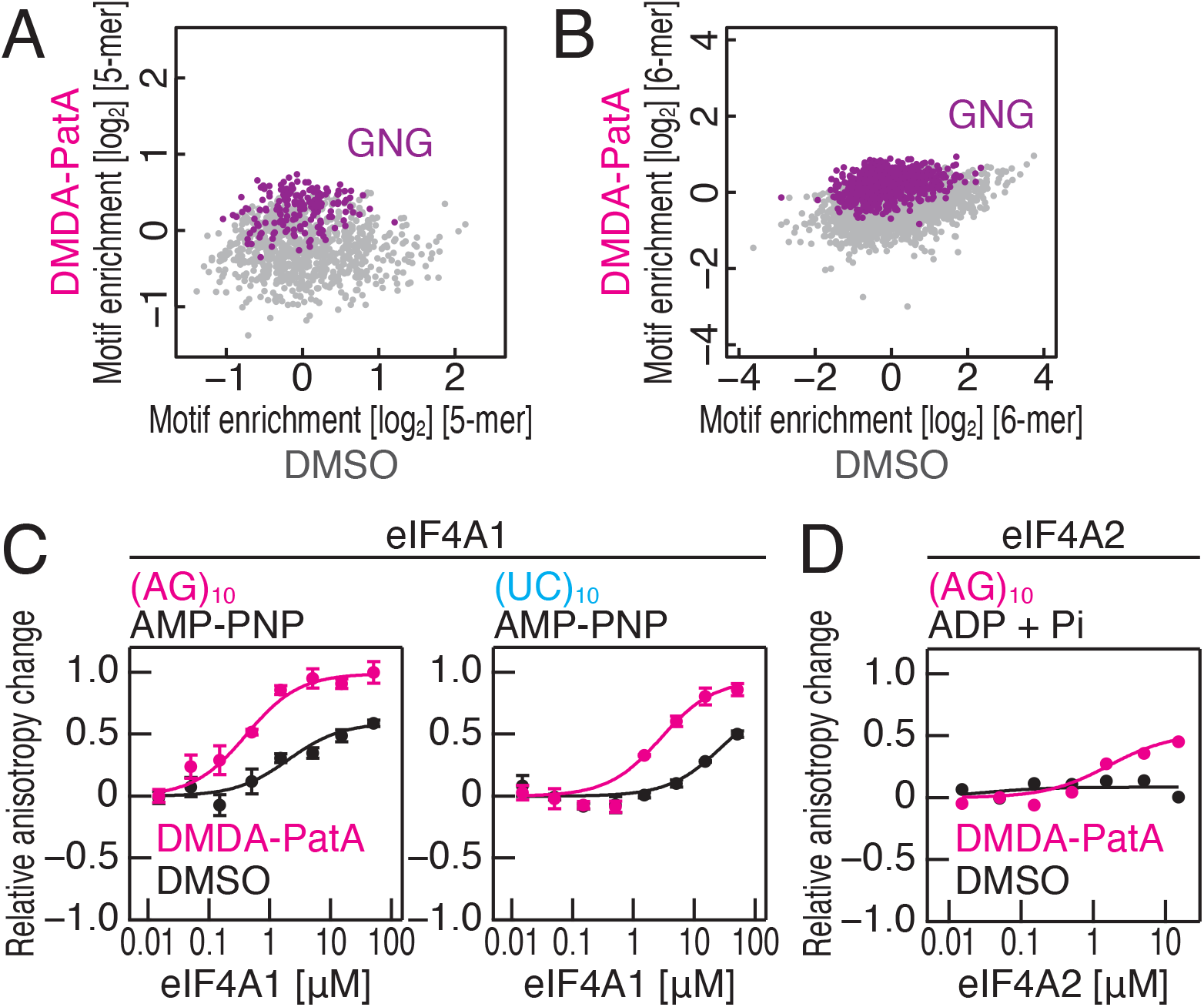
Validation of the results obtained by RNA Bind-n-Seq with a fluorescence polarization assay, related to Figure 1 and Table 1. (A and B) Comparison of enrichment of 5-mer motifs (A) or 6-mer motifs (B) with DMSO and those with DMDA-PatA. AMP-PNP and 15 pmol recombiannd eIF4A1 was included in the reaction. Motifs containing GNG are highlighted. (C and D) Fluorescence polarization assay for FAM-labeled RNAs along the titrated recombinant eIF4A1 (C) or eIF4A2 (D) with AMP-PNP (C) or ADP and Pi (D). The indicated RNA sequences at 10 nM were used with or without 50 μM DMDA-PatA. The data are presented as the mean (point) and s.d. (error) for replicates (n = 3).

**Figure S3.**
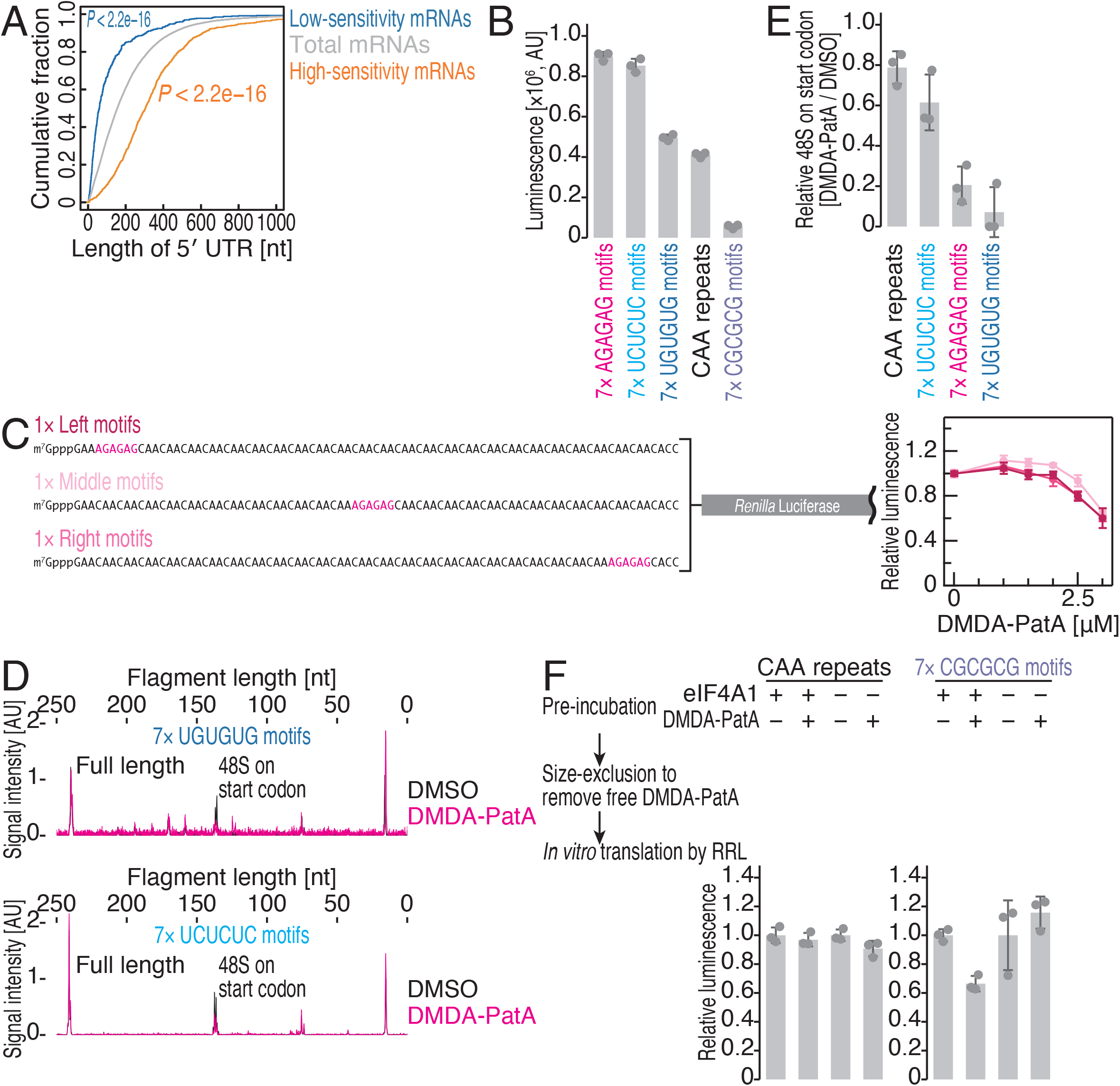
Characterization of the *in vitro* translation assay with DMDA-PatA, related to Figure 3. (A) Cumulative distribution of 5′ UTR length. DMDA-PatA low-sensitivity and high-sensitivity mRNAs (defined in Figure 1D) were compared to total mRNAs. The significance was calculated by the Mann‒Whitney *U* test. (B) Basal translation activities of the indicated reporter mRNAs in RRL. The data are presented as the mean (bar) and s.d. (error) for replicates (point, n = 3). (C) Schematic of reporter mRNAs with a 1× AGAGAG motif (left). These mRNAs were subjected to *in vitro* translation with RRL with the titration of DMDA-PatA. The data are presented as the mean (point) and s.d. (error) for replicates (n = 3). (D) Toeprinting assay to probe 48S ribosomes assembled on the start codons in the indicated reporter mRNAs with or without 10 μM DMDA-PatA. cDNA synthesized with FAM-labeled reverse transcription primers was analyzed by capillary electrophoresis. (E) The relative 48S formation on the start codons upon 10 μM DMDA-PatA treatment. The data are presented as the mean (bar) and s.d. (error) for replicates (point, n = 3). (F) *In vitro* translation of reporter mRNA (with 7× CGCGCG motifs and CAA repeats, at 181.8 nM) preincubated with recombinant eIF4A1 and DMDA-PatA. The size exclusion column was used to eliminate free DMDA-PatA. The data are presented as the mean (bar) and s.d. (error) for replicates (point, n = 3).

**Figure S4.**
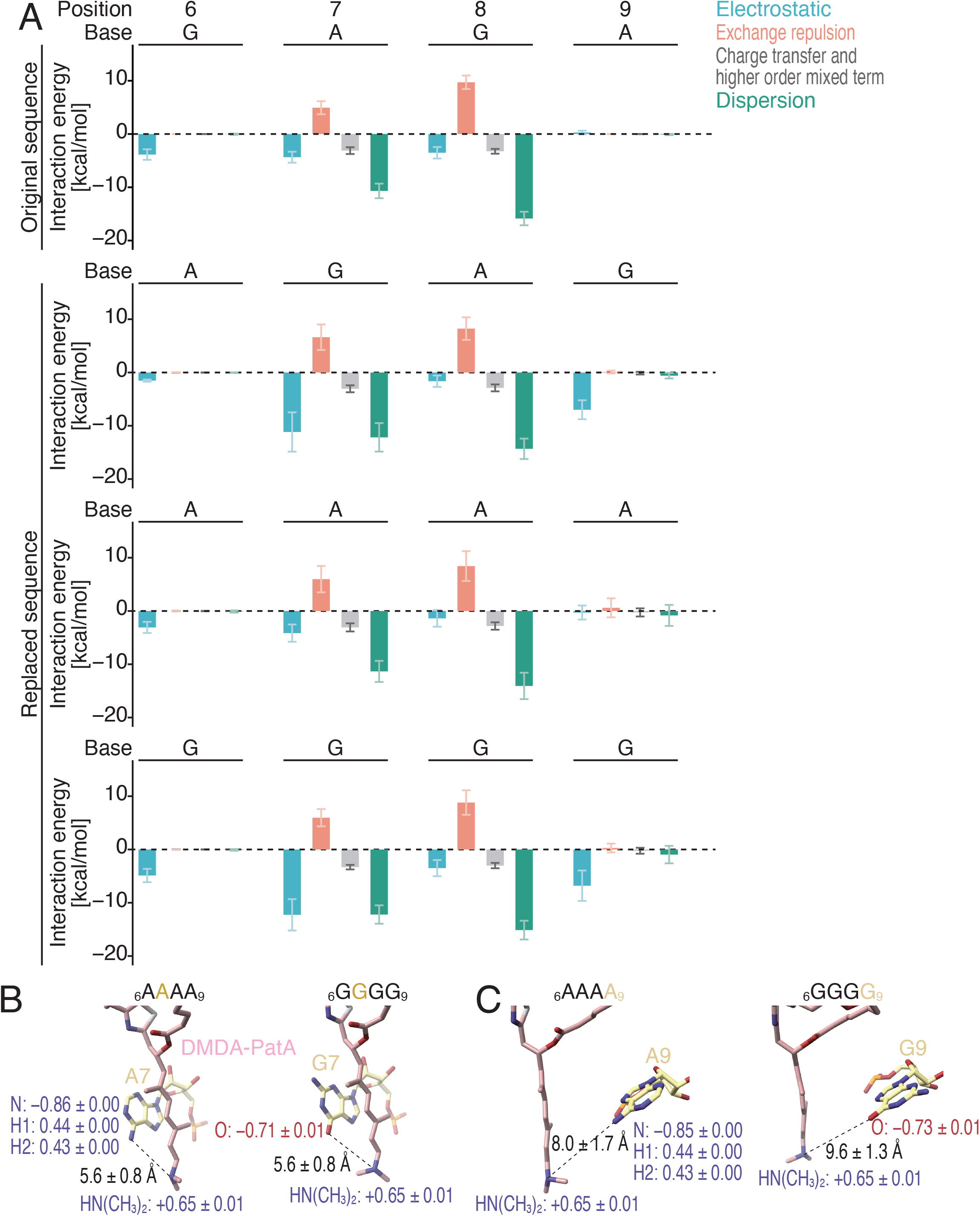
Characterization of the MD simulation and FMO calculation, related to Figure 4. (A) The interaction energy between DMDA-PatA and the indicated bases along the investigated complexes through FMO calculation. The interaction energy was decomposed to the indicated terms. The data present the mean (bar) and s.d. (error) for 10 complexes. (B and C) Representative structures obtained in MD simulation (at a time point of 100 ns) with _6_AAAA_9_ or _6_GGGG_9_. The net charges (*e*) on the indicated groups and the distances are shown (mean ± s.d. from 10 simulated structures).

**Figure S5.**
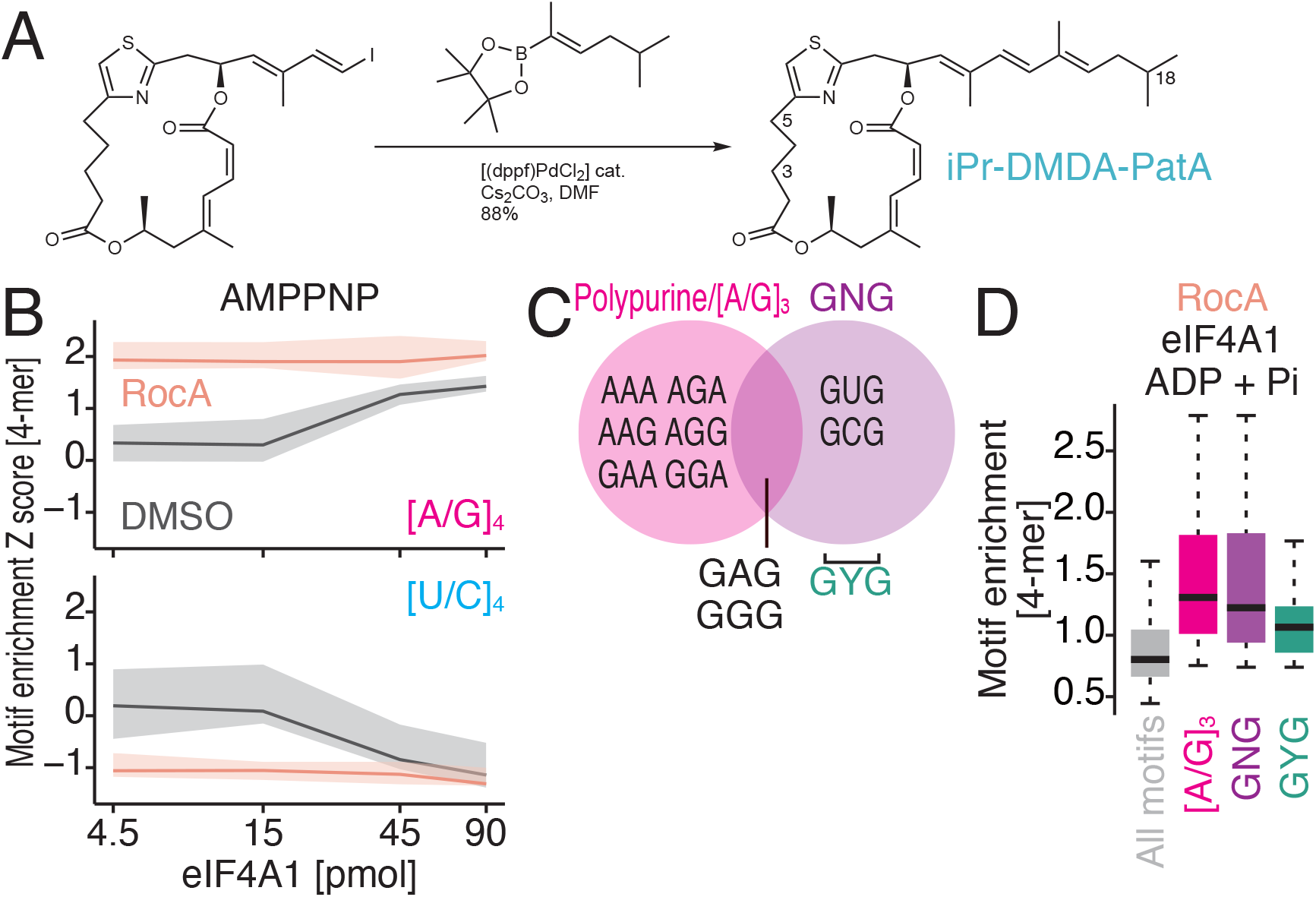
The differential sequence selectivity between RocA and DMDA-PatA on eIF4A1, related to Figure 5. (A) iPr-DMDA-PatA was readily attained by palladium-catalyzed Suzuki reaction of alkenyl iodide ^64,69^ with the known alkenylboronate ^67,68^. With this donor lacking the tertiary amine, the cross-coupling proceeded under sufficiently mild conditions that the isomerization-prone *Z,E*-configured dienoate embedded in the macrocyclic ring remained intact; no saponification of the lactone was observed either. This outcome is noteworthy, as the analogous attachment of the side chain with the –NMe_2_ terminus during the total syntheses of PatA as well as DMDA-PatA necessitated recourse to Stille reactions that were specifically optimized for applications to sensitive substrates ^79^. (B) Motif enrichment Z score along the titrated recombinant eIF4A1 in RNA Bind-n-Seq (with AMP-PNP) for the indicated 4-mer species with or without RocA. (C) Venn diagram to indicate the difference and commonality in motif specificities of RocA and DMDA-PatA. Y represents U or C. (D) Box plots for motif enrichments in RNA Bind-n-Seq (with ADP and Pi) on eIF4A1 with RocA in the indicated 4-mer species.

**Figure S6.**
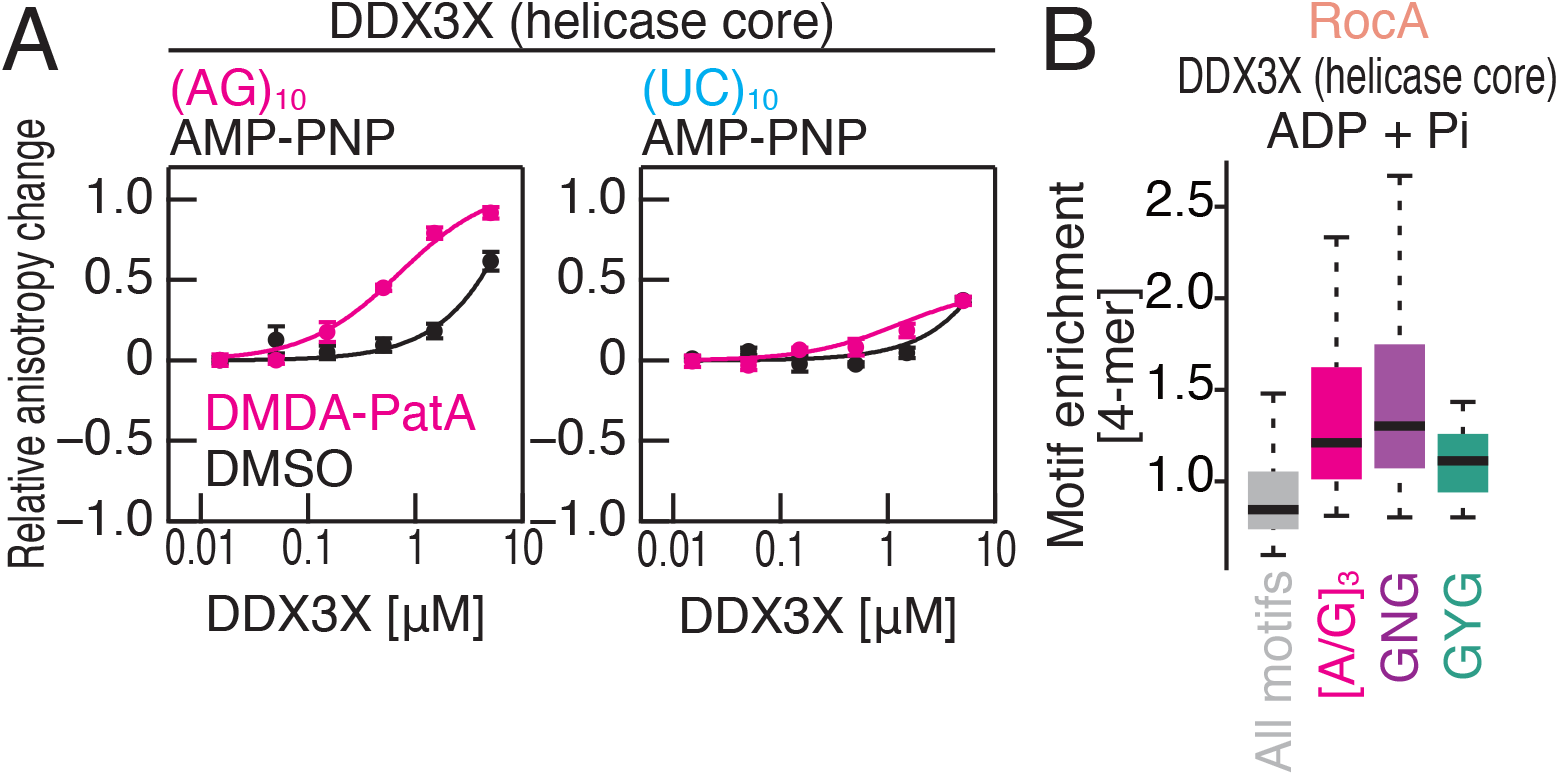
The differential sequence selectivity between RocA and DMDA-PatA on DDX3X, related to Figure 6 and Table 1. (A) Fluorescence polarization assay for FAM-labeled RNAs along the titrated recombinant DDX3X (helicase core) with AMP-PNP. The indicated RNA sequences at 10 nM were used with or without 50 μM DMDA-PatA. The data are presented as the mean (point) and s.d. (error) for replicates (n = 3). (B) Box plots of motif enrichments in RNA Bind-n-Seq (with ADP and Pi) on DDX3X (helicase core) with RocA in the indicated 4-mer species.

**Table S1. List of deep sequencing data used in this study, related to Figures 1-3, 5, and 6**.

Each data set is listed with library name, treatment, experiment batch, treatment, cell line, rRNA depletion method, replicate number, data type, reference, accession number, sequencer, and run mode.

**Table S2. List of FMODB IDs of FMO calculation results for each MD sampling structures, related to Figure 4**.

In box plots, the median (center line), upper/lower quartiles (box limits), and 1.5× interquartile range (whiskers) are shown.

## References

1. Valeur, E., and Jimonet, P. (2018). New modalities, technologies, and partnerships in probe and lead generation: enabling a mode-of-action centric paradigm. J. Med. Chem. 61, 9004–9029.

2. Shichino, Y., and Iwasaki, S. (2022). Compounds for selective translational inhibition. Curr. Opin. Chem. Biol. 69, 102158.

3. Vázquez-Laslop, N., and Mankin, A.S. (2018). Context-specific action of ribosomal antibiotics. Annu. Rev. Microbiol. 72, 185–207.

4. Atanasov, A.G., Zotchev, S.B., Dirsch, V.M., International Natural Product Sciences Taskforce, and Supuran, C.T. (2021). Natural products in drug discovery: advances and opportunities. Nat. Rev. Drug Discov. 20, 200–216.

5. Lin, J., Zhou, D., Steitz, T.A., Polikanov, Y.S., and Gagnon, M.G. (2018). Ribosome-targeting antibiotics: modes of action, mechanisms of resistance, and implications for drug design. Annu. Rev. Biochem. 87, 451–478.

6. Shen, L., and Pelletier, J. (2020). Selective targeting of the DEAD-box RNA helicase eukaryotic initiation factor (eIF) 4A by natural products. Nat. Prod. Rep. 37, 609–616.

7. Higa, T., Tanaka, J.-I., Tsukitani, Y., and Kikuchi, H. (1981). Hippuristanols, cytotoxic polyoxygenated steroids from the gorgonian *Isis hippuris*. Chem. Lett. 10, 1647–1650.

8. Bordeleau, M.E., Mori, A., Oberer, M., Lindqvist, L., Chard, L.S., Higa, T., Belsham, G.J., Wagner, G., Tanaka, J., and Pelletier, J. (2006). Functional characterization of IRESes by an inhibitor of the RNA helicase eIF4A. Nat. Chem. Biol. 2, 213–220.

9. Lindqvist, L., Oberer, M., Reibarkh, M., Cencic, R., Bordeleau, M.E., Vogt, E., Marintchev, A., Tanaka, J., Fagotto, F., Altmann, M., et al. (2008). Selective pharmacological targeting of a DEAD box RNA helicase. PLoS One 3, e1583.

10. Sun, Y., Atas, E., Lindqvist, L.M., Sonenberg, N., Pelletier, J., and Meller, A. (2014). Single-molecule kinetics of the eukaryotic initiation factor 4AI upon RNA unwinding. Structure 22, 941–948.

11. Steinberger, J., Shen, L., J Kiniry, S., Naineni, S.K., Cencic, R., Amiri, M., Aboushawareb, S.A.E., Chu, J., Maïga, R.I., Yachnin, B.J., et al. (2020). Identification and characterization of hippuristanol-resistant mutants reveals eIF4A1 dependencies within mRNA 5′ leader regions. Nucleic Acids Res. 48, 9521–9537.

12. King, M.L., Chiang, C.-C., Ling, H.-C., Fujita, E., Ochiai, M., and McPhail, A.T. (1982). *X*-Ray crystal structure of rocaglamide, a novel antileulemic 1*H*-cyclopenta[*b*]benzofuran from *Aglaia elliptifolia*. J. Chem. Soc. Chem. Commun. 1, 1150–1151.

13. Bordeleau, M.E., Robert, F., Gerard, B., Lindqvist, L., Chen, S.M., Wendel, H.G., Brem, B., Greger, H., Lowe, S.W., Porco, J.A., et al. (2008). Therapeutic suppression of translation initiation modulates chemosensitivity in a mouse lymphoma model. J. Clin. Invest. 118, 2651–2660.

14. Sadlish, H., Galicia-Vazquez, G., Paris, C.G., Aust, T., Bhullar, B., Chang, L., Helliwell, S.B., Hoepfner, D., Knapp, B., Riedl, R., et al. (2013). Evidence for a functionally relevant rocaglamide binding site on the eIF4A-RNA complex. ACS Chem. Biol. 8, 1519–1527.

15. Iwasaki, S., Floor, S.N., and Ingolia, N.T. (2016). Rocaglates convert DEAD-box protein eIF4A into a sequence-selective translational repressor. Nature 534, 558– 561.

16. Chu, J., Zhang, W., Cencic, R., Devine, W.G., Beglov, D., Henkel, T., Brown, L.E., Vajda, S., Porco, J.A., and Pelletier, J. (2019). Amidino-rocaglates: a potent class of eIF4A inhibitors. Cell Chem Biol 26, 1586–1593.e3.

17. Iwasaki, S., Iwasaki, W., Takahashi, M., Sakamoto, A., Watanabe, C., Shichino, Y., Floor, S.N., Fujiwara, K., Mito, M., Dodo, K., et al. (2019). The translation inhibitor rocaglamide targets a bimolecular cavity between eIF4A and polypurine RNA. Mol. Cell 73, 738–748.e9.

18. Chu, J., Zhang, W., Cencic, R., O’Connor, P.B.F., Robert, F., Devine, W.G., Selznick, A., Henkel, T., Merrick, W.C., Brown, L.E., et al. (2020). Rocaglates induce gain-of-function alterations to eIF4A and eIF4F. Cell Rep. 30, 2481– 2488.e5.

19. Northcote, P.T., Blunt, J.W., and Munro, M.H.G. (1991). Pateamine: a potent cytotoxin from the New Zealand Marine sponge, mycale sp. Tetrahedron Lett. 32, 6411–6414.

20. Bordeleau, M.E., Matthews, J., Wojnar, J.M., Lindqvist, L., Novac, O., Jankowsky, E., Sonenberg, N., Northcote, P., Teesdale-Spittle, P., and Pelletier, J. (2005). Stimulation of mammalian translation initiation factor eIF4A activity by a small molecule inhibitor of eukaryotic translation. Proc. Natl. Acad. Sci. U. S. A. 102, 10460–10465.

21. Low, W.K., Dang, Y., Schneider-Poetsch, T., Shi, Z., Choi, N.S., Merrick, W.C., Romo, D., and Liu, J.O. (2005). Inhibition of eukaryotic translation initiation by the marine natural product pateamine A. Mol. Cell 20, 709–722.

22. Bordeleau, M.E., Cencic, R., Lindqvist, L., Oberer, M., Northcote, P., Wagner, G., and Pelletier, J. (2006). RNA-mediated sequestration of the RNA helicase eIF4A by Pateamine A inhibits translation initiation. Chem. Biol. 13, 1287–1295.

23. Low, W.K., Dang, Y., Bhat, S., Romo, D., and Liu, J.O. (2007). Substrate-dependent targeting of eukaryotic translation initiation factor 4A by pateamine A: negation of domain-linker regulation of activity. Chem. Biol. 14, 715–727.

24. Low, W.-K., Li, J., Zhu, M., Kommaraju, S.S., Shah-Mittal, J., Hull, K., Liu, J.O., and Romo, D. (2014). Second-generation derivatives of the eukaryotic translation initiation inhibitor pateamine A targeting eIF4A as potential anticancer agents. Bioorg. Med. Chem. 22, 116–125.

25. Popa, A., Lebrigand, K., Barbry, P., and Waldmann, R. (2016). Pateamine A-sensitive ribosome profiling reveals the scope of translation in mouse embryonic stem cells. BMC Genomics 17, 52.

26. Chen, R., Zhu, M., Chaudhari, R.R., Robles, O., Chen, Y., Skillern, W., Qin, Q., Wierda, W.G., Zhang, S., Hull, K.G., et al. (2019). Creating novel translation inhibitors to target pro-survival proteins in chronic lymphocytic leukemia. Leukemia 33, 1663–1674.

27. Rust, M., Helfrich, E.J.N., Freeman, M.F., Nanudorn, P., Field, C.M., Rückert, C., Kündig, T., Page, M.J., Webb, V.L., Kalinowski, J., et al. (2020). A multiproducer microbiome generates chemical diversity in the marine sponge *Mycale hentscheli*. Proc. Natl. Acad. Sci. U. S. A. 117, 9508–9518.

28. Storey, M.A., Andreassend, S.K., Bracegirdle, J., Brown, A., Keyzers, R.A., Ackerley, D.F., Northcote, P.T., and Owen, J.G. (2020). Metagenomic exploration of the marine sponge *Mycale hentscheli* uncovers multiple polyketide-producing bacterial symbionts. MBio 11. 10.1128/mBio.02997-19.

29. Naineni, S.K., Liang, J., Hull, K., Cencic, R., Zhu, M., Northcote, P., Teesdale-Spittle, P., Romo, D., Nagar, B., and Pelletier, J. (2021). Functional mimicry revealed by the crystal structure of an eIF4A:RNA complex bound to the interfacial inhibitor, desmethyl pateamine A. Cell Chem Biol 28, 825–834.e6.

30. Santos, A.C., and Adkilen, P. (1932). The alkaloids of *Argemone mexicana*. J. Am. Chem. Soc. 54, 2923–2924.

31. Jiang, C., Tang, Y., Ding, L., Tan, R., Li, X., Lu, J., Jiang, J., Cui, Z., Tang, Z., Li, W., et al. (2019). Targeting the N terminus of eIF4AI for inhibition of its catalytic recycling. Cell Chem Biol 26, 1417–1426.e5.

32. Hinnebusch, A.G. (2014). The scanning mechanism of eukaryotic translation initiation. Annu. Rev. Biochem. 83, 779–812.

33. Brito Querido, J., Sokabe, M., Kraatz, S., Gordiyenko, Y., Skehel, J.M., Fraser, C.S., and Ramakrishnan, V. (2020). Structure of a human 48*S* translational initiation complex. Science 369, 1220–1227.

34. Chen, M., Asanuma, M., Takahashi, M., Shichino, Y., Mito, M., Fujiwara, K., Saito, H., Floor, S.N., Ingolia, N.T., Sodeoka, M., et al. (2021). Dual targeting of DDX3 and eIF4A by the translation inhibitor rocaglamide A. Cell Chem Biol 28, 475–486.e8.

35. Ingolia, N.T., Ghaemmaghami, S., Newman, J.R., and Weissman, J.S. (2009). Genome-wide analysis *in vivo* of translation with nucleotide resolution using ribosome profiling. Science 324, 218–223.

36. Iwasaki, S., and Ingolia, N.T. (2017). The growing toolbox for protein synthesis studies. Trends Biochem. Sci. 42, 612–624.

37. Romo, D., Choi, N.S., Li, S., Buchler, I., Shi, Z., and Liu, J.O. (2004). Evidence for separate binding and scaffolding domains in the immunosuppressive and antitumor marine natural product, pateamine a: design, synthesis, and activity studies leading to a potent simplified derivative. J. Am. Chem. Soc. 126, 10582–10588.

38. Iwasaki, S., Floor, S.N., and Ingolia, N.T. (2016). Rocaglates convert DEAD-box protein eIF4A into a sequence-selective translational repressor. Nature 534, 558– 561.

39. Liu, T.Y., Huang, H.H., Wheeler, D., Xu, Y., Wells, J.A., Song, Y.S., and Wiita, A.P. (2017). Time-resolved proteomics extends ribosome profiling-based measurements of protein synthesis dynamics. Cell Syst 4, 636–644.e9.

40. Chhipi-Shrestha, J.K., Schneider-Poetsch, T., Suzuki, T., Mito, M., Khan, K., Dohmae, N., Iwasaki, S., and Yoshida, M. (2022). Splicing modulators elicit global translational repression by condensate-prone proteins translated from introns. Cell Chemical Biology 29, 259–275.e10.

41. Naineni, S.K., Itoua Maïga, R., Cencic, R., Putnam, A.A., Amador, L.A., Rodriguez, A.D., Jankowsky, E., and Pelletier, J. (2020). A comparative study of small molecules targeting eIF4A. RNA 26, 541–549.

42. Lambert, N., Robertson, A., Jangi, M., McGeary, S., Sharp, P.A., and Burge, C.B. (2014). RNA Bind-n-Seq: quantitative assessment of the sequence and structural binding specificity of RNA binding proteins. Mol. Cell 54, 887–900.

43. Lambert, N.J., Robertson, A.D., and Burge, C.B. (2015). RNA Bind-n-Seq: measuring the binding affinity landscape of RNA-binding proteins. Methods Enzymol. 558, 465–493.

44. Linder, P., and Jankowsky, E. (2011). From unwinding to clamping — the DEAD box RNA helicase family. Nat. Rev. Mol. Cell Biol. 12, 505–516.

45. Weis, K., and Hondele, M. (2022). The role of DEAD-Box ATPases in gene expression and the regulation of RNA-protein condensates. Annu. Rev. Biochem. 91, 197–219.

46. Pestova, T.V., and Kolupaeva, V.G. (2002). The roles of individual eukaryotic translation initiation factors in ribosomal scanning and initiation codon selection. Genes Dev. 16, 2906–2922.

47. Dmitriev, S.E., Pisarev, A.V., Rubtsova, M.P., Dunaevsky, Y.E., and Shatsky, I.N. (2003). Conversion of 48S translation preinitiation complexes into 80S initiation complexes as revealed by toeprinting. FEBS Lett. 533, 99–104.

48. Shirokikh, N.E., Alkalaeva, E.Z., Vassilenko, K.S., Afonina, Z.A., Alekhina, O.M., Kisselev, L.L., and Spirin, A.S. (2010). Quantitative analysis of ribosome-mRNA complexes at different translation stages. Nucleic Acids Res. 38, e15.

49. Chen, M., Kumakura, N., Saito, H., Muller, R., Nishimoto, M., Mito, M., Gan, P., Ingolia, N.T., Shirasu, K., Ito, T., et al. (2023). A parasitic fungus employs mutated eIF4A to survive on rocaglate-synthesizing *Aglaia* plants. Elife 12, e81302.

50. Fedorov, D., and Kitaura, K. (2009). Fragment Molecular Orbital Method Practical Applications to Large Molecular Systems (Taylor & Francis Group).

51. Fedorov, D.G., Nagata, T., and Kitaura, K. (2012). Exploring chemistry with the fragment molecular orbital method. Phys. Chem. Chem. Phys. 14, 7562–7577.

52. Tanaka, S., Mochizuki, Y., Komeiji, Y., Okiyama, Y., and Fukuzawa, K. (2014). Electron-correlated fragment-molecular-orbital calculations for biomolecular and nano systems. Phys. Chem. Chem. Phys. 16, 10310–10344.

53. Mochizuki, Y., Tanaka, S., and Fukuzawa, K. (2021). Recent Advances of the Fragment Molecular Orbital Method: Enhanced Performance and Applicability (Springer Nature Singapore).

54. Handa, Y., Okuwaki, K., Kawashima, Y., Hatada, R., Mochizuki, Y., Komeiji, Y., Tanaka, S., Furuishi, T., Yonemochi, E., Honma, T., et al. (2023). Prediction of binding pose and affinity of SARS-CoV-2 Main Protease and repositioned drugs by combining docking, molecular dynamics, and fragment molecular orbital calculations. ChemRxiv. 10.26434/chemrxiv-2023-hrsvj.

55. Amari, S., Aizawa, M., Zhang, J., Fukuzawa, K., Mochizuki, Y., Iwasawa, Y., Nakata, K., Chuman, H., and Nakano, T. (2006). VISCANA: visualized cluster analysis of protein-ligand interaction based on the ab initio fragment molecular orbital method for virtual ligand screening. J. Chem. Inf. Model. 46, 221–230.

56. Tsukamoto, T., Kato, K., Kato, A., Nakano, T., Mochizuki, Y., and Fukuzawa, K. (2015). Implementation of pair interaction energy decomposition analysis and its applications to protein-ligand systems. J. Comput. Chem. Jpn. 14, 1–9.

57. Mullard, A. (2017). Small molecules against RNA targets attract big backers. Nat. Rev. Drug Discov. 16, 813–815.

58. Garber, K. (2023). Drugging RNA. Nat. Biotechnol. 41, 745–749.

59. Khaperskyy, D.A., Emara, M.M., Johnston, B.P., Anderson, P., Hatchette, T.F., and McCormick, C. (2014). Influenza a virus host shutoff disables antiviral stress-induced translation arrest. PLoS Pathog. 10, e1004217.

60. González-Almela, E., Sanz, M.A., García-Moreno, M., Northcote, P., Pelletier, J., and Carrasco, L. (2015). Differential action of pateamine A on translation of genomic and subgenomic mRNAs from Sindbis virus. Virology 484, 41–50.

61. Ziehr, B., Lenarcic, E., Cecil, C., and Moorman, N.J. (2016). The eIF4AIII RNA helicase is a critical determinant of human cytomegalovirus replication. Virology 489, 194–201.

62. Slaine, P.D., Kleer, M., Smith, N.K., Khaperskyy, D.A., and McCormick, C. (2017). Stress granule-inducing eukaryotic translation initiation factor 4A inhibitors block influenza A virus replication. Viruses 9, 388.

63. Romo, D., Rzasa, R.M., Shea, H.A., Park, K., Langenhan, J.M., Sun, L., Akhiezer, A., and Liu, J.O. (1998). Total Synthesis and Immunosuppressive Activity of (−)-Pateamine A and Related Compounds: Implementation of a β-Lactam-Based Macrocyclization. J. Am. Chem. Soc. 120, 12237–12254.

64. Zhuo, C.-X., and Fürstner, A. (2018). Catalysis-based total syntheses of pateamine A and DMDA-Pat A. J. Am. Chem. Soc. 140, 10514–10523.

65. Guan, W., Michael, A.K., McIntosh, M.L., Koren-Selfridge, L., Scott, J.P., and Clark, T.B. (2014). Stereoselective formation of trisubstituted vinyl boronate esters by the acid-mediated elimination of α-hydroxyboronate esters. J. Org. Chem. 79, 7199–7204.

66. McIntosh, M.L., Moore, C.M., and Clark, T.B. (2010). Copper-catalyzed diboration of ketones: facile synthesis of tertiary alpha-hydroxyboronate esters. Org. Lett. 12, 1996–1999.

67. Xu, S., Geng, P., Li, Y., Liu, G., Zhang, L., Guo, Y., and Huang, Z. (2021). Pincer Iron Hydride Complexes for Alkene Isomerization: Catalytic Approach to Trisubstituted (Z)-Alkenyl Boronates. ACS Catal. 11, 10138–10147.

68. Sanchez, A., and Maimone, T.J. (2022). Taming shapeshifting anions: total synthesis of ocellatusone C. J. Am. Chem. Soc. 144, 7594–7599.

69. Zhuo, C.-X., and Fürstner, A. (2016). Concise synthesis of a pateamine A analogue with in vivo anticancer activity based on an iron-catalyzed pyrone ring opening/cross-coupling. Angew. Chem. Int. Ed Engl. 55, 6051–6056.

70. Mito, M., Mishima, Y., and Iwasaki, S. (2020). Protocol for disome profiling to survey ribosome collision in humans and zebrafish. STAR Protoc 1, 100168.

71. Kashiwagi, K., Shichino, Y., Osaki, T., Sakamoto, A., Nishimoto, M., Takahashi, M., Mito, M., Weber, F., Ikeuchi, Y., Iwasaki, S., et al. (2021). eIF2B-capturing viral protein NSs suppresses the integrated stress response. Nat. Commun. 12, 1– 12.

72. Langmead, B., and Salzberg, S.L. (2012). Fast gapped-read alignment with Bowtie 2. Nat. Methods 9, 357–359.

73. Anders, S., and Huber, W. (2010). Differential expression analysis for sequence count data. Genome Biol. 11, R106.

74. Dobin, A., Davis, C.A., Schlesinger, F., Drenkow, J., Zaleski, C., Jha, S., Batut, P., Chaisson, M., and Gingeras, T.R. (2013). STAR: ultrafast universal RNA-seq aligner. Bioinformatics 29, 15–21.

75. Chen, S., Zhou, Y., Chen, Y., and Gu, J. (2018). fastp: an ultra-fast all-in-one FASTQ preprocessor. Bioinformatics 34, i884–i890.

76. Mochizuki, Y., Nakano, T., Koikegami, S., Tanimori, S., Abe, Y., Nagashima, U., and Kitaura, K. (2004). A parallelized integral-direct second-order Møller–Plesset perturbation theory method with a fragment molecular orbital scheme. Theor. Chem. Acc. 112, 442–452.

77. Mochizuki, Y., Koikegami, S., Nakano, T., Amari, S., and Kitaura, K. (2004). Large scale MP2 calculations with fragment molecular orbital scheme. Chem. Phys. Lett. 396, 473–479.

78. Mochizuki, Y., Yamashita, K., Murase, T., Nakano, T., Fukuzawa, K., Takematsu, K., Watanabe, H., and Tanaka, S. (2008). Large scale FMO-MP2 calculations on a massively parallel-vector computer. Chem. Phys. Lett. 457, 396–403.

79. Fürstner, A., Funel, J.-A., Tremblay, M., Bouchez, L.C., Nevado, C., Waser, M., Ackerstaff, J., and Stimson, C.C. (2008). A versatile protocol for Stille-Migita cross coupling reactions. Chem. Commun., 2873–2875.

